# A genome-wide association study of serum proteins reveals shared loci with common diseases

**DOI:** 10.1101/2021.07.02.450858

**Authors:** Alexander Gudjonsson, Valborg Gudmundsdottir, Gisli T Axelsson, Elias F Gudmundsson, Brynjolfur G Jonsson, Lenore J Launer, John R Lamb, Lori L Jennings, Thor Aspelund, Valur Emilsson, Vilmundur Gudnason

**Author notes:** These authors contributed equally as joint-first authors (Alexander Gudjonsson, Valborg Gudmundsdottir). These authors contributed equally as joint-senior authors (Valur Emilsson, Vilmundur Gudnason).

## Abstract

With the growing number of genetic association studies, the genotype-phenotype atlas has become increasingly more complex, yet the functional consequences of most disease associated alleles is not understood. The measurement of protein level variation in solid tissues and biofluids integrated with genetic variants offers a path to deeper functional insights. Here we present a large-scale proteogenomic study in 5,368 individuals, revealing 4,113 independent associations between genetic variants and 2,099 serum proteins, of which 37% are previously unreported. The majority of both *cis*- and *trans*-acting genetic signals are unique for a single protein, although our results also highlight numerous highly pleiotropic genetic effects on protein levels and demonstrate that a protein’s genetic association profile reflects certain characteristics of the protein, including its location in protein networks, tissue specificity and intolerance to loss of function mutations. Integrating protein measurements with deep phenotyping of the cohort, we observe substantial enrichment of phenotype associations for serum proteins regulated by established GWAS loci, and offer new insights into the interplay between genetics, serum protein levels and complex disease.

## Main

The identification of causal genes underlying common diseases has the potential to reveal novel therapeutic targets and provide readouts to monitor disease risk. Genome-wide association studies (GWAS) have identified thousands of genetic variants conferring risk of disease, however, the highly polygenic architecture of most common disorders^1^ implies that the genetic component of common diseases is largely mediated through complex biological networks^2,3^. Identifying the causal mediators of mapped phenotype-associated genetic variation remains a largely unresolved challenge as majority of such variants reside in non-coding regulatory regions of the genome^4^. In fact, disease risk loci are enriched in regions of active chromatin involved in gene regulation^5,6^. Thus, the integration of intermediate molecular traits like mRNA^7^ or proteins^8–12^ with genetics and phenotypic information may aid the identification of causal candidates and functional consequences. Furthermore, the phenotypic pleiotropy observed at many loci^13^ calls for a better understanding of the chain of events that are introduced by disease associated variants. Genetic perturbations may for instance drive molecular cascades through regulatory networks^8^, most of which have not yet been fully mapped, or as a consequence of their phenotypic effects. Such downstream effects of genetic variants can be reflected in the molecular pleiotropy observed at some genetic loci, which may have important implications for therapeutic discovery including for estimating potential side effects^14^. For instance, many GWAS risk loci for complex diseases regulate multiple proteins in *cis* and *trans*, which often cluster in the same co-regulatory network modules^8^. Through the serum proteome we can gain a broad and well-defined description of the downstream effects of genetic variants, and their complex relationship with disease relevant traits.

The human plasma proteome consists of proteins that are secreted or shed into the circulation, either to carry out their function there or to mediate cross-tissue communications^15^. Proteins may also leak from tissues, for example as a result of tissue damage^15^. It has been noted that a large subset of *cis*-to-*trans* serum protein pairs (i.e. proteins that are regulated by the same genetic variant in *cis* or *trans*, respectively) have tissue specific expression but often involving distinct organ systems^8^, indicating that proteins in circulation may originate from virtually any tissue in the body. This suggests that system level coordination is facilitated to a considerable degree by proteins in blood, which if perturbed may mediate common disease^16^. These observations, together with the accessibility of blood compared to other tissues, make circulating proteins an attractive source for identifying molecular signatures of disease in large cohorts.

Recent technological advances now allow for high-throughput quantification of circulating proteins, which has resulted in the first large-scale studies^8–12^ of protein quantitative trait loci (pQTLs) as recently reviewed^17^. Here, we present a large-scale proteogenomic study revealing thousands of independent genetic loci affecting a substantial proportion of the serum proteome, highlighting widespread pleiotropic effects of disease-associated genetic variation on serum protein levels. While our previous work reported associations to a restricted set of loci^8^, this is the first comprehensive GWAS for this number of serum proteins. A systematic integrative analysis furthermore demonstrates extensive associations between serum proteins and phenotypes that are regulated by the same genetic signals, adding further support to the therapeutic target and biomarker potential among proteins regulated by established GWAS risk variants.

## Results

### Identification of cis and trans acting protein quantitative trait loci (pQTLs)

We performed a GWAS of 4,782 serum proteins encoded by 4,135 unique human genes in the population-based AGES cohort of elderly Icelanders (n = 5,368, Table S1), measured by the slow-off rate modified aptamer (SOMAmer) platform as previously described^8,18^. On average the genomic inflation factor was low (mean λ = 1.045, sd = 0.033) and of the 7,506,463 genetic variants included in the analysis (Fig. S1), 269,637 variants exhibited study-wide significant associations (P < 5×10^−8^/4,782 SOMAmers = 1.046×10^−11^) with 2,112 unique proteins, dubbed protein quantitative trait loci (pQTLs). In a conditional analysis, we identified 4,113 study-wide significant associations between 2,087 independent genetic signals in 799 loci (defined as genetic signals within 300kb of each other) and 2,099 unique proteins (Fig. 1A-C, Tables S2-S4). Here we defined a genetic signal as a set of genetic variants in linkage disequilibrium (LD) that were associated with one or more proteins. For each associated protein, a genetic signal has a lead variant, defined as the genetic variant that is most confidently associated with the protein, i.e. with the lowest P-value (see Methods for details). Among the 4,113 independent associations, those in *cis* (signal lead variant within 300kb of the protein-encoding gene boundaries, n = 1,429) tended to have larger effect sizes than those in *trans* (signal lead variant >300kb from the protein-encoding gene boundaries, n = 2,684) (Fig. S2A). We found that almost half (977/2,099 = 47%) of all proteins with any independent genetic associations had more than one signal (Fig. 1B). Of those, 579 proteins (59%) had more than one independent signal within the same locus (Fig. S2B) and 697 proteins (71%) had signals in distinct locations in the genome. The protein with the largest number of associated loci was TENM3 (10 loci), followed by NOG (9 loci), GRAMD1C and TMCC3 (7 loci each).

**Fig. 1.**
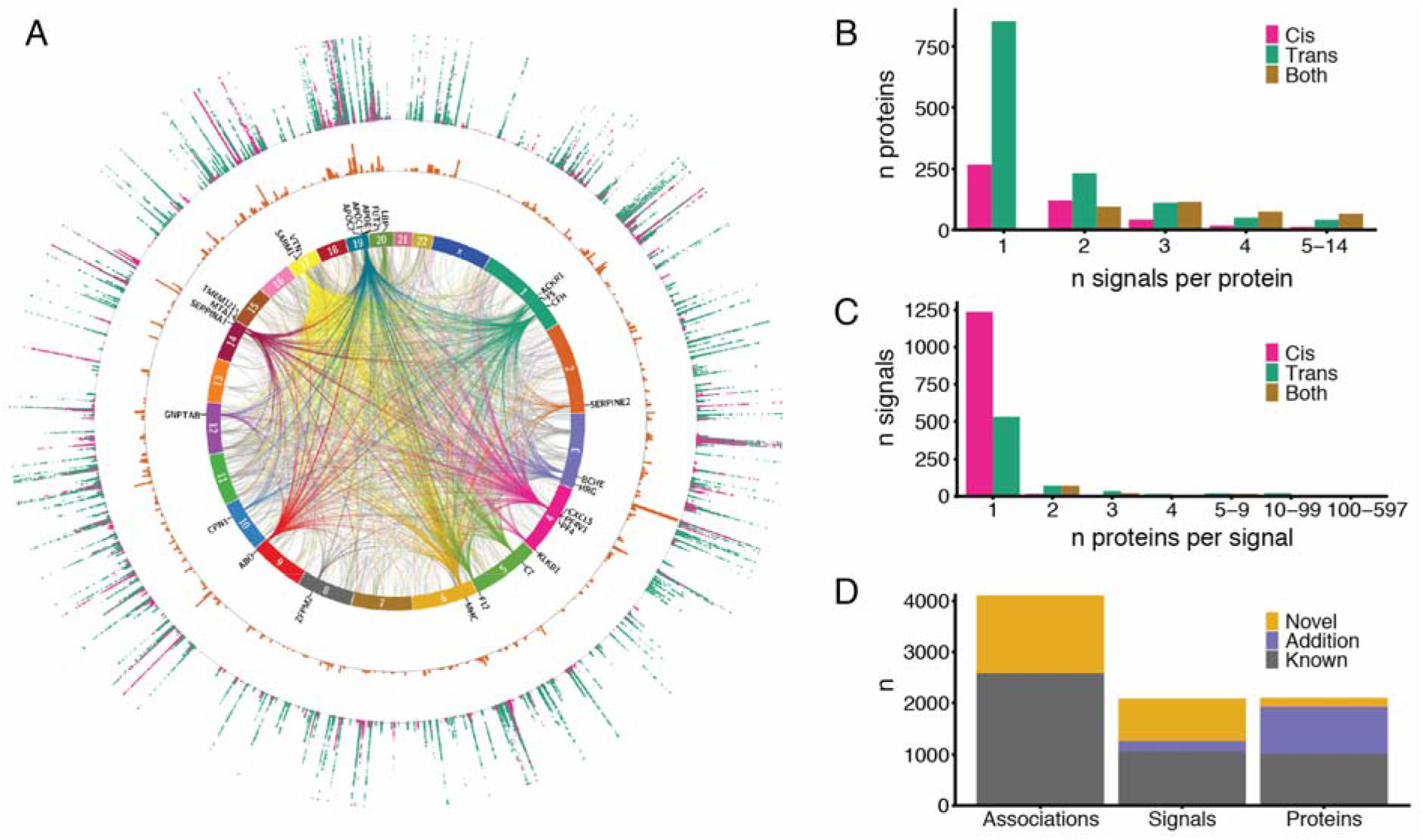
A) Circos plot showing every study-wide significant variant-protein association from the protein GWAS (n = 5,368). The innermost layer shows links between independent signals and *trans* gene locations of associated proteins. *Trans* hotspots are colored by the chromosome they originate from. The second layer states the nearest genes to these *trans* hotspots. The third layer is a histogram of the distribution of the independent signals, where each bar represents the number of independent signals within 300kb from each other, values ranging from 1 to 38. The outermost layer is a Manhattan plot for all proteins, P-values ranging from 1×10^−11^ to 1×10^−300^ (capped), colored by *cis* (pink) or *trans* (green). B) Barplot showing number of proteins, binned by the number of associated independent signals, colored by *cis* (pink), *trans* (green) or both (mustard). C) Barplot showing number of independent signals, binned by the number of associated proteins, colored by *cis* (pink), *trans* (green) or both (mustard). D) Barplot showing the number of novel associations compared to similar large-scale genotype-protein association studies.

The majority of genetic signals were only associated with a single protein (Fig.1C), or 98% of *cis* signals and 73% of *trans* signals, and can as such be considered specific for the given protein based on a recently proposed classification of *trans*-pQTLs^11^. Furthermore, we have previously shown that proteins regulated in *trans* by the same genetic variant often cluster in the same coregulatory networks, sharing functionality and a disease relationship, although they may often differ in tissue origin^8^. However, as in previous studies^8–11^, we identified numerous hotspots of *trans* protein associations, or more specifically 35 independent signals that were associated with 10 or more proteins each at a study-wide significant threshold (Fig. 1A,C). The largest of these *trans* hotspots represents the variant rs704, a missense variant within the Vitronectin (*VTN*) gene, which was associated with 598 proteins. Many of these *trans* hotspots are well established as such, including the *VTN*, *ABO*, *APOE, CFH* and *BCHE* loci^8–11^. Other notable *trans* hotspots included for instance variants in or near the Lipopolysaccharide Binding Protein (*LBP)* and Metastasis-Associated 1 (*MTA1)* genes. *LBP* is involved in the innate immune response to bacterial infections and *MTA1* encodes a transcriptional coregulator upregulated in numerous cancer types and associated with cancer progression^19^. Of the 35 *trans* hotspots, 14 also affected protein levels encoded by proximal genes, thus acting in *cis* as well (Table S3).

In contrast to the *trans* acting hotspots, we also observed genetic regions with high density of independent signals, each of which were not necessarily associated with many proteins. One such region stood out in particular on chromosome 3 (Fig. 1A), where 30 independent signals were observed for a total of 55 proteins within a 300kb window (Fig. S3A), of which six proteins (ADIPOQ, AHSG, DNAJB11, FETUB, HRG and KNG1) were regulated in *cis*. Further analysis of this region demonstrated a sparse LD structure (Fig. S3A), allowing for this high density of independent signals, and revealing a subcluster of 15 genetic signals affecting 32 proteins in various constellations (Fig. S3B), that were enriched for Toll Like Receptor 7/8 cascade (FDR = 4.8×10^−3^) and MAP kinase activation (FDR = 4.8×10^−3^).

To define what proportion of the pQTLs identified in the present study can be considered novel, we compared all study-wide significant pQTLs with previously reported pQTL studies (Table S5), including the recent exome array analysis of the AGES cohort^20^. Of the 4,113 independent associations detected in the current study, 1,527 (37.1%) are considered novel based on this comparison (Supplementary Note 1, Fig. 1E, Fig. S4). Of the 2,087 independent genetic signals, 821 (39.3%) are novel, in the sense that they have not been reported to associate with any protein, and we find new protein associations for 206 known signals. Out of the 2,099 proteins, 172 (8.2%) had no previously reported genetic associations in the comparison and we identified new genetic associations for additional 911 proteins.

We evaluated how well independent pQTLs reported by the INTERVAL study^9^ (n = 3,301) replicated in our results and found 75.6% to be both directionally consistent and nominally significant (P < 0.05) (Supplementary Note 2, Fig. S5-S6). This proportion furthermore increased to 93.9% when the *NLRP12* locus was excluded, a reported *trans* hotspot that did not replicate in the AGES cohort (Supplementary Note 2, Fig. S5-S6). This locus has in fact been identified as platform specific in a recent study^21^ and was suggested to be related to white blood cell lysis during sample handling. We similarly performed a lookup of the independent study-wide significant associations identified in the current study in the INTERVAL study summary statistics (Supplementary Note 2, Fig. S7). Of 2,716 associations with information in the INTERVAL study we find that 94.1% are directionally consistent and 82.0% were both directionally consistent and nominally significant (P < 0.05). Of 668 associations defined as novel in our study (Supplementary Note 1) and with information available in the INTERVAL study, we again find a very high directional consistency between the two studies, or 89.8% of associations, and 62.9% are both directionally consistent and nominally significant (P < 0.05) in the smaller INTERVAL study.

Finally, with more individuals genotyped we revisited the GWAS of the serum protein co-regulatory network^8^, now represented by the first two eigenproteins of each module, and find that almost all the network modules are under strong genetic control (Supplementary Note 3).

### Characterization of proteins by genetic association profiles

Taking advantage of the broad coverage of the protein measurements in our study, to determine which protein characteristics can provide additional insights into the observed differences in genetic profiles for the measured proteins we compared characteristics such as tissue-enhanced gene^22^ and protein^23^ expression and protein localization^22^ for proteins with genetic signals to those without any detected genetic effect. Moreover, we analyzed loss-of-function (LoF) intolerance^24^ and hub status in two types of protein networks, i.e. the InWeb protein-protein interaction (PPI) network^25^ and the serum protein co-regulatory network^8^, but pathogenicity of DNA sequence variation and hub status of proteins in biological networks are well-known features used to study the extent of selection pressure in molecular evolution^26,27^. We find that proteins with study-wide significant genetic associations, specifically those acting in *cis*, are generally more likely to have tissue-specific gene and protein expression and are more often secreted compared to those with no detected genetic signals (Fig. 2A, Tables S6-S7). These results may indicate that that *cis*-pQTLs in serum to some extent mirror the regulation of protein secretion from solid tissues, whereas the serum level of proteins without *cis*-pQTLs may mainly be affected by other mechanisms. By contrast, proteins with *trans* only signals are enriched among transmembrane proteins (Fig. 2A, Tables S6-S7). Furthermore, we find that proteins with *cis* signals generally have lower LoF intolerance, that is they are more tolerant to deleterious mutations, and they tend to have lower hub status in both PPI and co-regulatory networks, indicating a more peripheral position of *cis* regulated proteins in protein networks (Fig. 2B, Tables S6-S7). Similarly, larger genetic effects on protein levels are negatively correlated with LoF intolerance and hub status in both the PPI and co-regulatory networks (Fig. S8). This suggests that selective pressure may to some extent explain the lack of pQTLs for proteins that are encoded by housekeeping genes, are network hubs and are intolerant to LoF mutations.

**Fig. 2.**
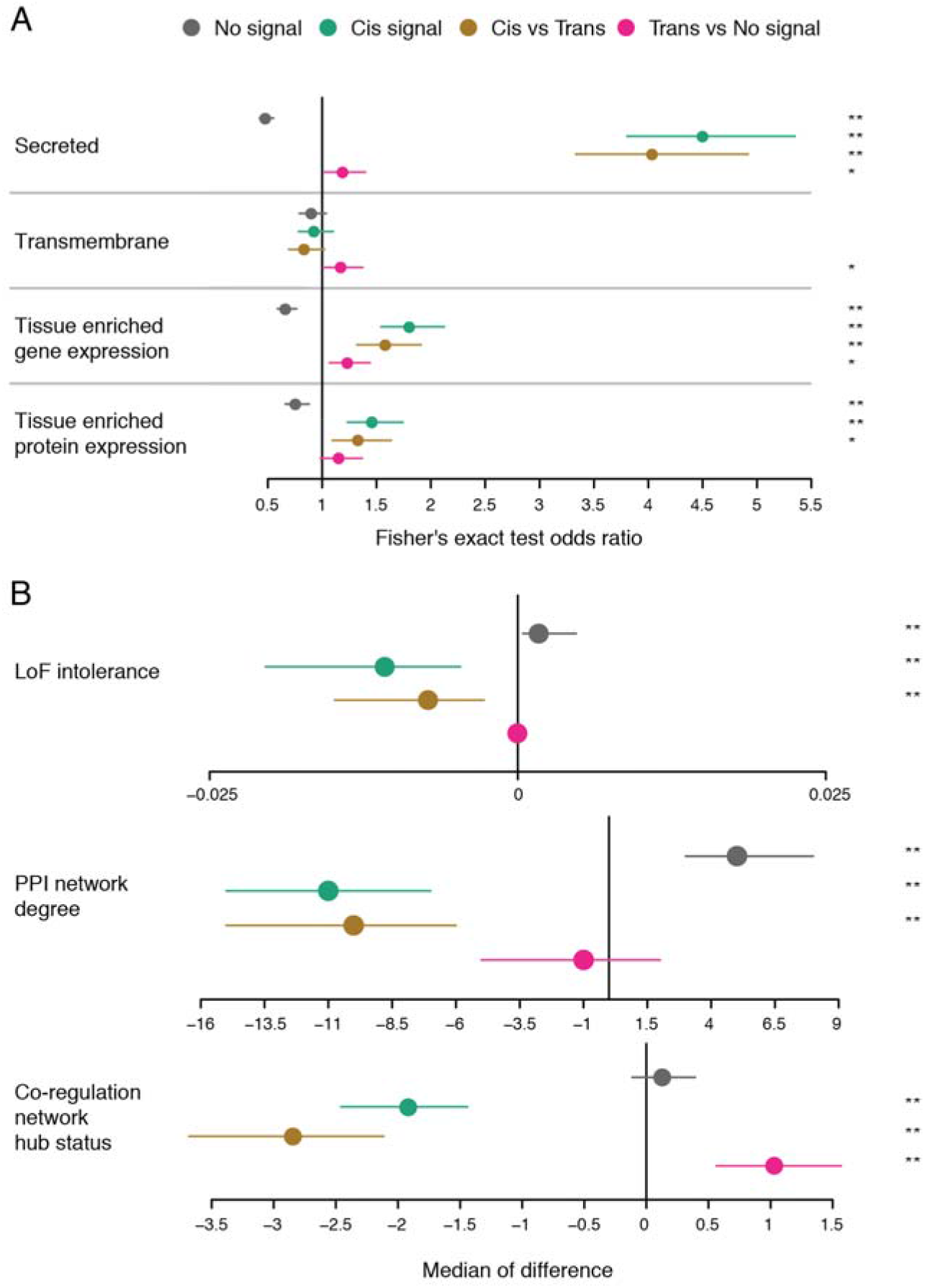
Enrichment analysis estimates and 95% confidence intervals comparing characteristics between proteins classified by types of genetic association signals. See main text for definitions. A) Fisher’s exact test for comparing classifications. B) Wilcoxon’s rank sum test for comparing classifications with continuous traits. The estimate and confidence interval represents the median of the difference between values from the two classes. The stars on the right indicate statistical significance; * p < 0.05, ** p < 0.001.

Proteins with *trans* acting signals had higher hub status in the co-regulatory network compared to those proteins having no genetic signals (Fig. 2B). However, *trans* signals were not associated with hub status in the PPI network or influenced by LoF intolerance (Fig. 2B). Complementing this observation, we find that hub proteins in co-regulatory networks are generally connected to more proteins through the same genetic variants (Fig. S8). As the co-regulatory network is derived from protein correlations, these results highlight how its structure is to some extent shaped by genetic variants affecting multiple proteins, the majority of which are *trans* regulated^8^ (Supplementary Note 3). These results elucidate key differences between the PPI and the serum protein co-regulatory networks, i.e. while hubs in both types of networks are depleted for *cis*-pQTLs, only those in the co-regulatory network were more likely *trans*-regulated proteins.

### Colocalization of pQTLs with GWAS risk loci

Genetic effects on serum proteins may offer novel insights into mechanisms underlying the genetics of common disease and relevant traits. Therefore, we examined the overlap between pQTLs and GWAS loci. We obtained GWAS summary statistics for 81 diseases and clinical traits (Table S8) and identified all genome-wide significant (P < 5×10^−8^) GWAS loci overlapping with a study-wide significant pQTL from our results. Of note, the number of significant loci for each of the tested phenotypes is highly dependent on the original study size (Fig. S9). GWAS signals for different phenotypes were considered to belong to the same locus if the lead variants were within 300kb of each other. By this criteria, 1,335 GWAS loci for 76 phenotypes were found to be in the vicinity of a study-wide significant pQTL and were tested for colocalization. Of those, 218 GWAS loci (associated with 69 phenotypes) had high support (PP4>0.8) for colocalization with 1,045 proteins (Fig. 3, Tables S9-S10). Additionally, medium support (0.5<P4<=0.8) was found for colocalization between 171 proteins and 84 loci associated with 49 phenotypes (Fig. 3, Tables S9-S10). Of the 799 loci associated with protein levels, 216 (27.4%) colocalized with at least one GWAS phenotype and 1,083 (51%) of the 2,112 proteins with a study-wide significant pQTL. We found 91% (69/76) of the phenotypes tested to have a genetic signal colocalizing with at least one protein, with an average of 9 (11%) colocalized loci per trait (Fig. S10). GWAS loci with *cis*-pQTLs were more likely to colocalize (medium or high support) with any protein than those without (22.3% vs 10.4%, Fisher’s exact test P = 7.5×10^−8^). For a given phenotype, we observed that its associated loci involved a median of 17 serum proteins (Fig. S11). Thus, even a limited proportion of associated loci for a given phenotype generally associates with numerous proteins in serum and consequently implicate multiple affected molecular pathways. To account for multiple independent signals in a given locus, we additionally ran a conditional colocalization analysis for loci that had more than one independent signal per protein, thus including 549 GWAS loci that overlapped with pQTLs for 546 proteins. Here we observed 178 instances of colocalization with medium or high support, of which 51 (involving 19 loci, 14 phenotypes and 40 proteins) were not captured in the initial colocalization analysis (Tables S11-S12).

**Fig. 3.**
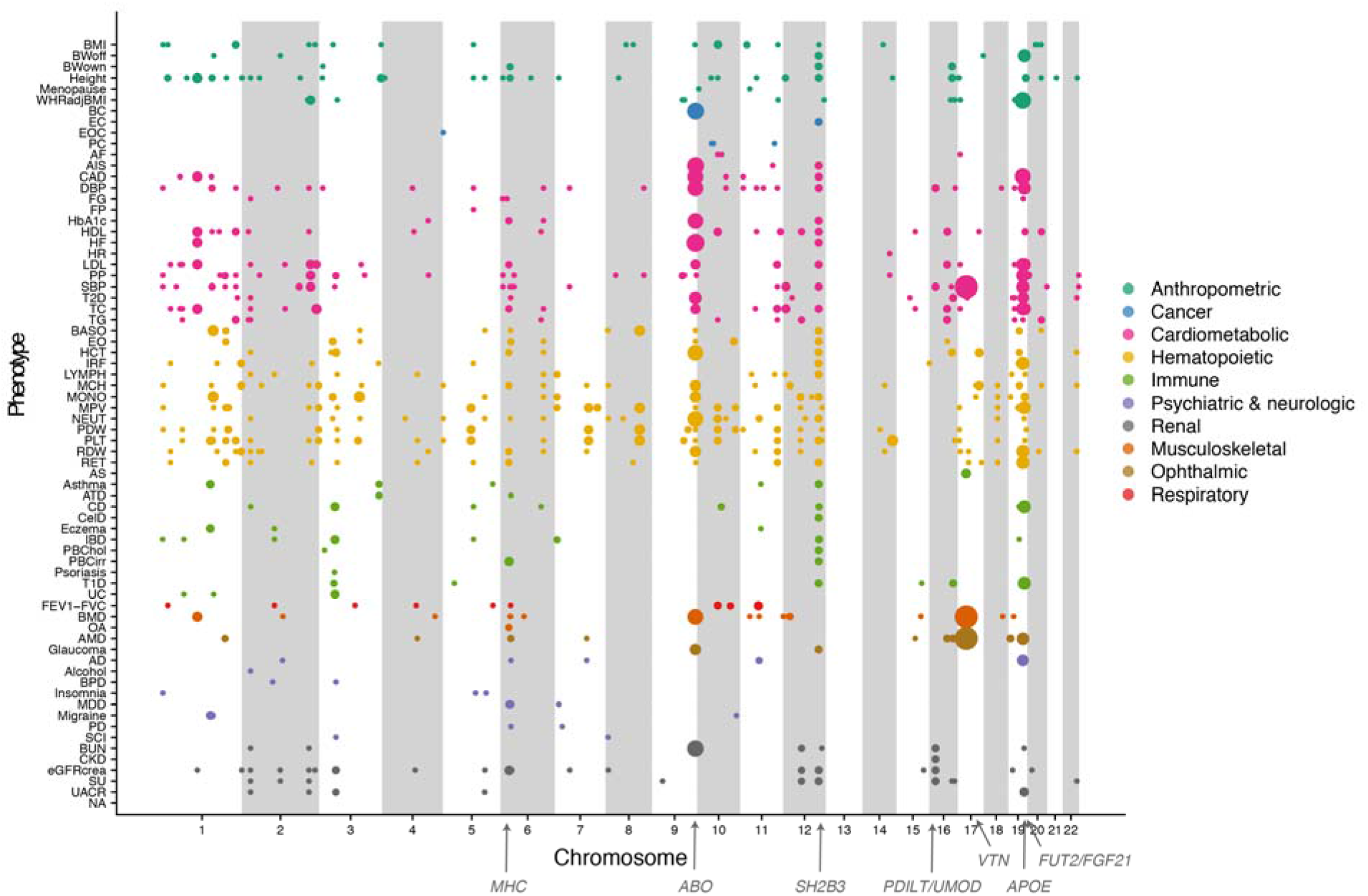
Overivew of colocalization between protein and phenotype associations across the genome. Each dot represents a genetic locus (genomic location on x-axis) that is associated with a phenotype (y-axis), where the dots size indicates the number of colocalized proteins (coloc PP4>0.5). Phenotype abbreviations are available from Table S8.

Colocalized *cis*-acting pQTLs can point to causal genes at GWAS loci. We found 237 and 49 trait-locus-*cis-*protein combinations with high or medium support, respectively. For 102 of 203 (50.2%) unique pairs of GWAS lead variants and colocalized *cis*-pQTLs, the protein was different than that encoded by the nearest gene to the GWAS lead variant (Table S10). For example, a GWAS signal for waist-to-hip ratio in the gene *LRRC36*, colocalizes with a pQTL for the serum levels of Agouti-related protein encoded by a nearby gene, *AGRP* (Fig. S12), a neuropeptide that increases appetite and decreases metabolism^28^. A related example involves two loci associated with BMI, located 5Mb apart on chromosome 20, both of which colocalize with serum levels of the Agouti signaling protein (ASIP) (Fig. S13), known to promote obesity via the melanocortin receptor (MC4R)^29^. These two associations are 2.2Mb and 7.6Mb upstream of the *ASIP* gene, respectively, however the colocalization with serum levels of ASIP suggest this may in fact be the causal candidate mediating their effects. Among neurological phenotypes, colocalized *cis*-pQTL examples include a GWAS signal for bipolar disorder on chromosome 2, which colocalizes with the serum levels of the protein encoded by *LMAN2L* (Fig. S14), and a signal for major depression disorder on chromosome 7 colocalizing with TMEM106B (Fig. S14), adding support for these being the causal genes at these loci, both of which are also the nearest gene to the GWAS lead variant.

We observed several highly pleiotropic loci, where multiple phenotype signals colocalized with multiple protein signals (Fig. 4A). In fact, among the high (PP4>0.8) and medium confidence (PP4>0.5) colocalization results, the number of associated proteins per GWAS locus was positively correlated with the number of associated phenotypes (Spearman’s rho = 0.50, P = 9.9×10^−17^). These pleiotropic loci included for example the *ABO* locus, best known for its role in determining the ABO blood groups, which was found to harbor eight independent protein signals within a 28 kb region (chr 9, 136,127,268-136,155,127) (Table S4), where pQTLs for 63 proteins colocalized with 17 phenotypes, predominantly cardiometabolic and hematopoietic (Fig. 4A, Table S10). The complex genetic architecture at this locus gives rise to a wide range of downstream consequences, as indicated by the distinct sets of proteins associated with each independent genetic signal defined here and consistent with previous reports^10^, and most traits associated with the locus are affected by more than one of those signals. The 63 proteins in the *ABO* locus were enriched for gene ontology terms and pathways such as “transmembrane signaling receptor activity” (FDR = 2.7×10^−6^), “regulation of cell migration” (FDR = 2.5×10^−4^) and “Hippo-Merlin signaling dysregulation” (FDR = 1.2×10^−3^).

**Fig. 4.**
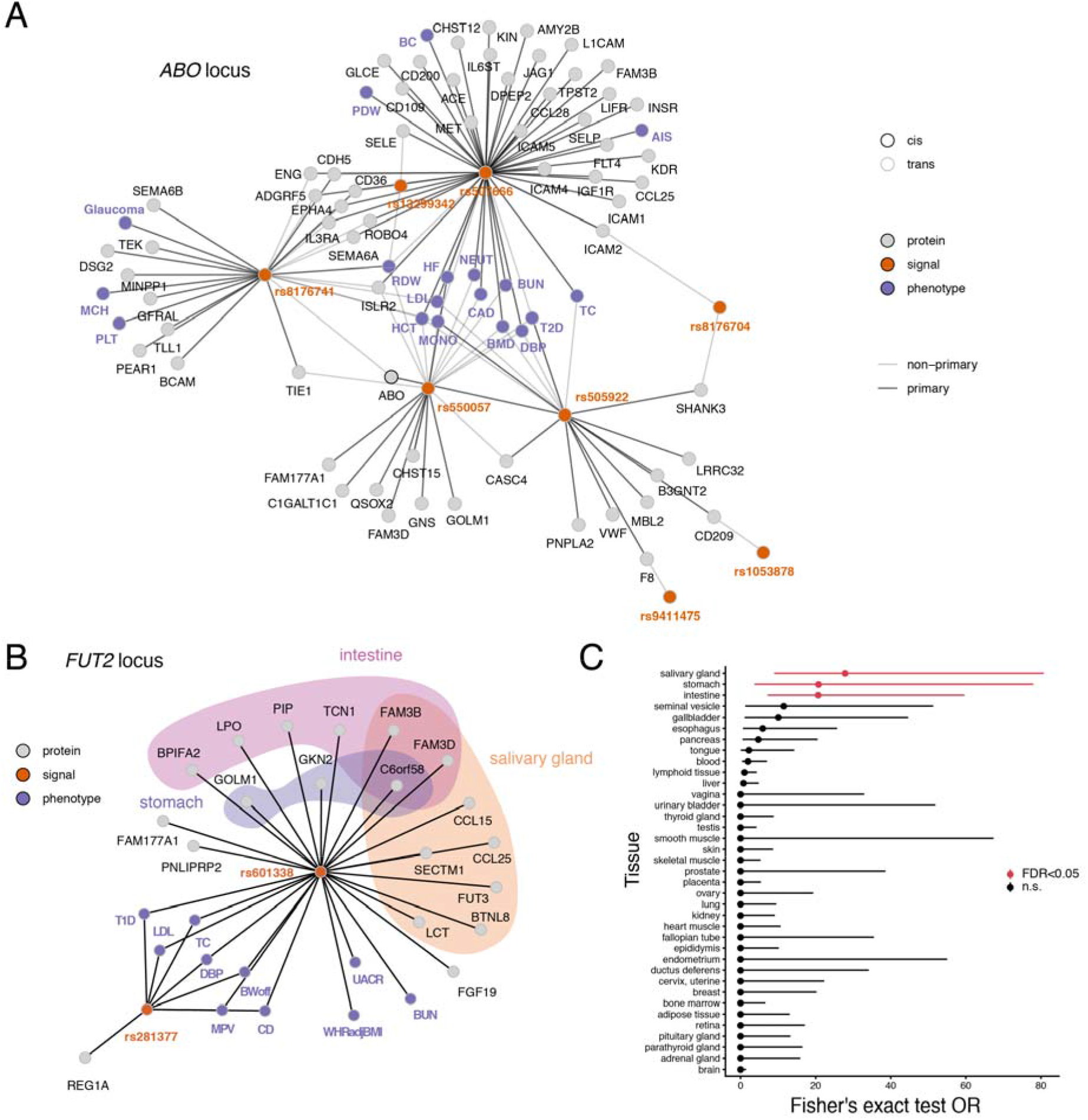
A) An overview of independent genome-wide significant genetic signals (orange nodes), annotated by the SNP with the strongest protein association, at the *ABO* locus (chr 9, 136,127,268 – 136,155,127) and their links to proteins (grey nodes) and phenotypes (purple nodes). Edges between genetic signals and proteins indicate primary (dark edges) and secondary (light edges) independent signals from the conditional analysis. Edges between genetic signals and traits indicate that any of the lead pQTL SNPs within that signal reaches P<<5×10^−8^ in GWAS summary statistics for the given trait, and the primary signal is assigned for the trait based on the lowest P-value. B) An overview of the independent genome-wide significant genetic signals (orange nodes), annotated by the SNP with the strongest protein association, at the *FUT2* locus (chr 19, 49,206,108 – 49,252,151) and their links to proteins (grey nodes) and the phenotypes they colocalize with (purple nodes). The background color indicates tissue-elevated expression in salivary gland, intestine or stomach. C) Enrichment (Fisher’s exact test) of tissue-elevated expression among the 19 proteins regulated by the *FUT2* locus where Benjamini-Hochberg FDR<0.05 is considered significant (red). Phenotype abbreviations are available from Table S8.

Another example of a pleiotropic locus is a 46 kb window (chr 19, 49,206,108-49,252,151), harboring variants adjacent to or within *FUT2* that are associated with diverse traits (Fig. 4B, Table S10), including immune (Crohn’s disease and type 1 diabetes), anthropometric (waist-to-hip ratio and offspring birth weight), cardiometabolic (blood pressure, LDL and total cholesterol) and renal (BUN and UACR). *FUT2* encodes for fucosyltransferase-2 that synthesizes the H antigen in body fluids and the intestinal mucosa, while a nearby gene, *FGF21,* is an important metabolic regulator^30^, acting for example through its effects on sugar intake^31^. We find that the genetic signals for 10 phenotypes in this region colocalize with 19 proteins that are collectively enriched for elevated gene expression^22^ in the intestine (FDR = 1.4×10^−6^), salivary gland (FDR = 1.7×10^−6^) and stomach (FDR = 8.9×10^−3^) (Fig. 4B-C) and include proteins involved in carbohydrate digestion (LCT), taste perception (LPO, PIP) or humoral immunity (CCL25). The proteins regulated by this locus thus suggest downtream effects across different parts of the gastrointestinal tract. The shared genetic architecture of immune disorders has been well documented in the literature and is mirrored in multiple colocalized pQTLs shared between various immune diseases (Fig. S15). In particular the *SH2B3* locus on chromosome 12 stands out in this regard, with GWAS signals for seven immune disorders colocalizing with three *trans*-regulated proteins (THPO, ICAM2, CXCL11), all involved in positive regulation of immune system processes (GO:0002684).

In some cases we observed more than one colocalized *trans*-pQTLs converging on the same protein for a given phenotype. For example, HDL-associations in the *LIPC* (chromosome 15) and *APOB* (chromosome 2) loci both colocalized with the serum levels of the sodium-coupled transporter SLC5A8 (Fig. S16), involved in the transport of monocarboxylates such as lactate and short-chain fatty acids. Similarly, variants in the *GALNT2* (chromosome 1) and *GCKR* loci (chromosome 2) both regulate the serum levels of NRP1, colocalizing with GWAS signals for triglyceride levels (Fig. S17). A more extreme example is a network of 12 loci with GWAS signals for platelet counts that colocalize with serum levels of 24 proteins (Fig. S18). These proteins include noggin (NOG) and cochlin (COCH), colocalizing with platelet count signals in five and four loci, respectively.

### Associations of proteins with phenotypes in the AGES cohort

Taking advantage of the deep phenotyping of the AGES cohort, we examined direct associations between colocalized proteins and 37 phenotypes that were measured in the AGES cohort (Table S13). For a quarter (10/37) of the phenotypes tested we observed a significant enrichment of phenotype associations among the sets of colocalized proteins compared to randomly sampled proteins (Fig. 5, Fig. S19, Table S14), demonstrating more generally that GWAS loci for complex phenotypes regulate serum proteins that themselves are often directly associated to the phenotype itself. At a more relaxed genome-wide significant (P<5×10^−8^) threshold for pQTLs, the proportion of phenotypes with significant enrichment of protein associations increased to 45% (18/40 phenotypes, Fig. S20), likely due to an increase in statistical power with more colocalized proteins per phenotype at this threshold and indicating that more associations between proteins regulated by GWAS-loci and the respective phenotypes can be expected to be identified as sample sizes for proteogenomic studies increase. Among the diseases and clinical traits with the strongest enrichment for direct protein-trait associations, we found age-related macular degeneration (14% of colocalized proteins associated compared to an average of 7% for random proteins, P<0.001), total cholesterol (67% vs 35% for random, P<0.001), Alzheimer’s disease (21% vs 1% for random, P=0.001) and type 2 diabetes (60% vs 40% for random, P=0.017). In some cases, this enrichment was driven by proteins regulated from a few *trans* loci, as evident by the loss of significance when the analysis was repeated without pleiotropic loci regulating five or more proteins, leaving on average 17 proteins per trait (Fig. 5, Table S14). This was particularly evident for Alzheimer’s disease, where the enrichment was entirely driven by the associations of proteins regulated by the *APOE* locus (Table S13). In other cases, the removal of proteins regulated by pleiotropic loci resulted in an enhanced enrichment of phenotype associations, such as for HbA1c, mean platelet volume and diastolic blood pressure (Fig. S19, Table S14).

**Fig. 5.**
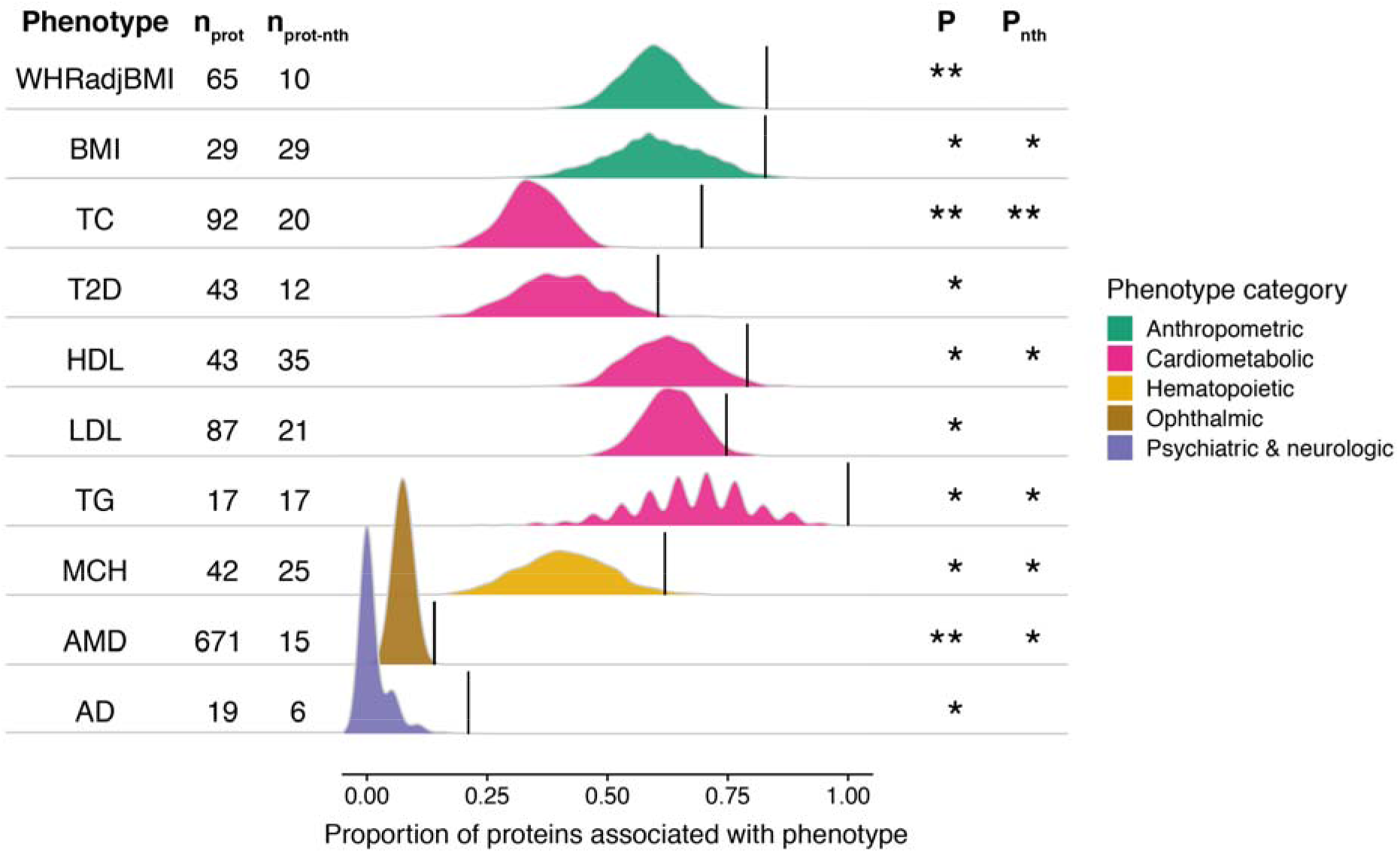
Ridgeline plot illustrating for each GWAS phenotype the proportion of colocalized proteins that were significantly (FDR<0.05) associated with the same trait in AGES (n = 5,457) (black lines) compared to 1000 randomly sampled sets of proteins of the same size (density curves), here showing only those with empirical P<0.05, see full results in Fig. S19. The number of colocalized proteins for each trait are provided on the left-hand side, along with the number of proteins remaining after the removal of proteins originating from loci with 5 or more colocalized proteins from the analysis, annotated as no transhotspots (nth). Empirical p-values for significant enrichment of trait-associations are denoted as such: *P < 0.05, **P < 0.001. WHRadjBMI, waist-to-hip ratio adjusted for BMI; TC, total cholesterol; T2D, type 2 diabetes; HDL, high-density lipoprotein cholesterol; LDL, low-density lipoprotein cholesterol; TG, triglycerides; MCH, mean corpuscular hemoglobin; AMD, age-related macular degeneration; AD Alzheimer’s disease.

By evaluating each individual locus separately, we identified six loci with significant phenotype-association enrichment among its linked proteins that colocalized with GWAS signals for the respective phenotype, thus demonstrating specific examples of genetic variants whose molecular and phenotypic consequences are linked within the same cohort (Table S15). Here the *APOE* locus stood out in terms of number of enriched phenotypes, with its regulated proteins being enriched for associations with Alzheimer’s disease, age-related macular degeneration, numerous cardiometabolic traits including coronary artery disease. The 641 proteins regulated by the *VTN* locus on chromosome 17 were also enriched for associations with AMD. The *PSRC1*-*CELSR2*-*SORT1* locus, best known for its associations with coronary artery disease and cholesterol levels, showed enrichment for protein associations with bone mineral density. Proteins regulated by the *ABO* locus on chromosome 9 and the *UGT* gene family cluster on chromosome 8 were enriched for associations with total cholesterol and finally the proteins regulated by the *ZFPM2* locus on chromosome 8 were enriched for associations with basophil counts. These genetic loci thus demonstrate specific examples whose molecular and phenotypic consequences are linked within the same cohort.

Other examples of colocalized proteins showing significant associations with the respective phenotype include the inhibin beta subunit B (INHBB) protein, which has a *cis*-pQTL on chromosome 2 and a *trans*-signal on chromosome 12, near the *INHBC* gene that encodes another subunit of the same protein complex, both of which colocalize with GWAS signals for estimated glomerular filtration rate (eGFR), a marker of renal function (Fig. 6A-C). The INHBB protein itself is associated with eGFR in the AGES cohort in a directionally consistent manner (Fig. 6C-D). Thus, the associations of these genetic variants affecting different components of the same protein complex together with the consistent association between the protein itself and eGFR indicate a possible role for the inhibin/activin proteins in renal function. Another example is the colocalization between a GWAS signal for type 2 diabetes with the missense lead variant rs738409 in the *PNPLA3* gene, a well established locus for non-alcoholic fatty liver disease^32^, and a *trans*-pQTL for ADP Ribosylation Factor Interacting Protein 2 (ARFIP2) (Fig. 6E), which is strongly downregulated in type 2 diabetes patients in AGES (Fig. 6F)^18^. These observations raise a number of new questions, for example how a missense variant in *PNPLA3* leads to a change in the circulating levels of ARFIP2, if ARFIP2 provides some sort of readout of *PNPLA3* function and finally how ARFIP2 relates to type 2 diabetes, i.e. if it mediates any of the risk associated with this locus or if it is merely a bystander. Thus more generally, the novel links between genetic loci, proteins and disease risk observed here can be used to derive new hypotheses for further studies.

**Fig. 6.**
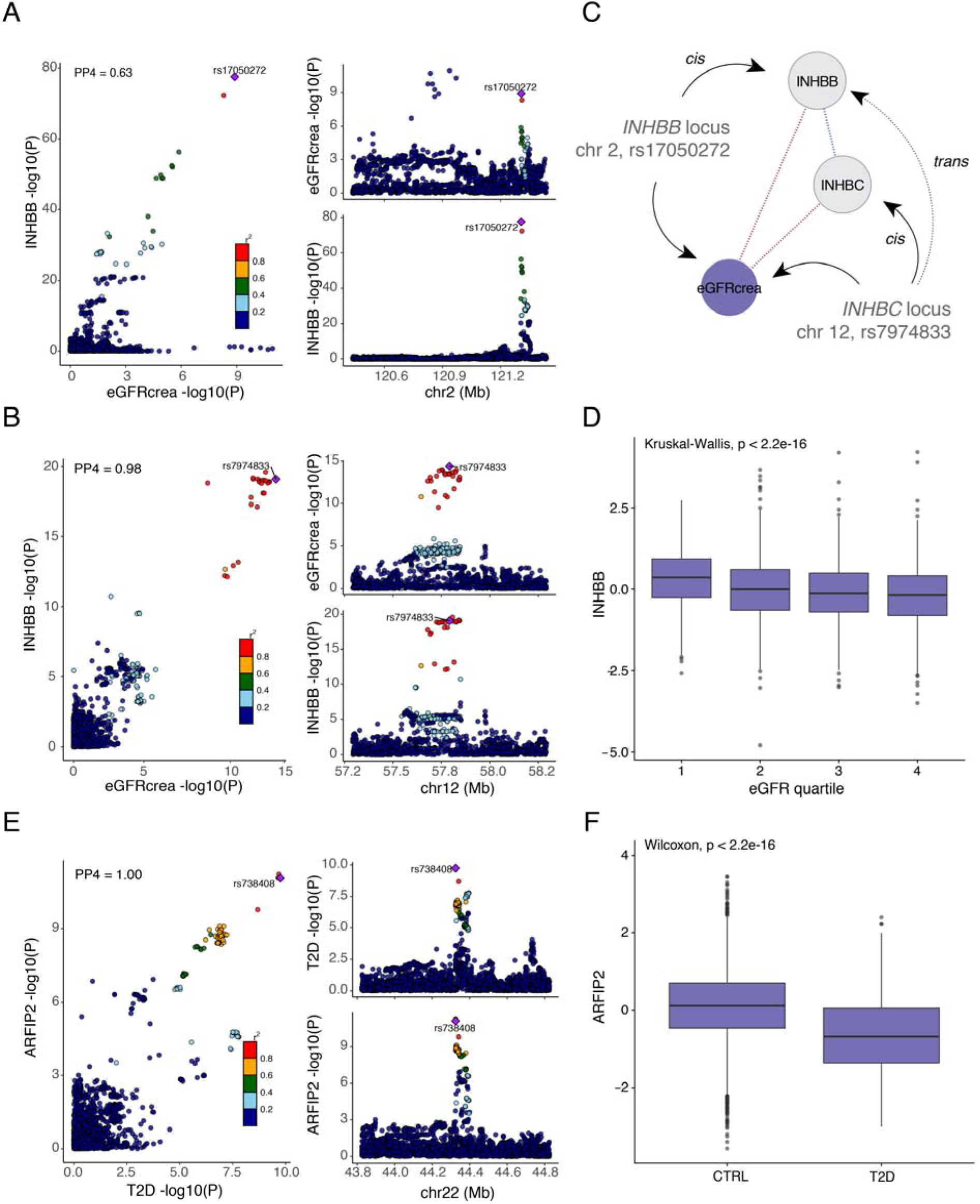
A-B, Colocalization between GWAS signals for eGFR and INHBB at A) the *INHBB* locus on chromosome 2 and B) the *INHBC* locus on chromosome 12. C) A schematic diagram showing the convergence of genetic effects on serum levels of INHBB at the *INHBB* locus in *cis* and *INHBC* locus in *trans*. Variants in the *INHBC* locus furthermore affect INHBC serum levels in *cis,* albeit not reaching study-wide significance (P = 8.5×10^−8^). Serum levels of INHBB and INHBC are positively correlated (Pearson’s r = 0.32, P = 3.4×10^−130^), while both are negatively associated with eGFR (beta = −4.52, SE = 0.23, P = 1.3×10^−82^ and beta = −2.62, SE = 0.22, P = 5.4×10^−32^, respectively).

## Discussion

In this work, we present the largest genome-wide association study of serum protein levels to date in terms of protein coverage, and demonstrate a substantial increase in existing knowledge as regards the number of significant genetic associations to proteins in circulation. We furthermore provide a systematic evaluation of protein-phenotype associations in the context of established risk loci for numerous diseases and clinical traits.

The current study expands on our previous work^8^ by increasing the number of genetic variants included in the analysis (from *cis*-regions only to a genome-wide analysis), thus increasing the search space, but also enhancing statistical power for identifying genetic associations by increasing the sample size in genetic analyses from 3,219 previously to 5,368 participants in the current study. Here, we identified study-wide significant genetic signals for half of the measured proteins and up to 16 independent genetic signals for a given protein. Thus, as for any other traits, the expected number of genetic associations for serum proteins can only be expected to increase with larger sample sizes, as has been demonstrated for CRP^33^. Large-scale meta-analyses across cohorts and biobanks will with time provide a more complete understanding of the genetic regulation of individual circulating proteins and their networks, including the effect of variability between different tissues on serum protein levels. The majority of c*is* and *trans* acting pQTLs detected in serum and plasma can be readily replicated across different populations, as shown in the current study, and different proteomic platforms^8,9,17,21^. However, a recent cross-platform comparison has shown that a subset of pQTLs are platform-specific and may in some cases represent epitope effects or other technical factors^21^. Thus, meta-analyses across platforms will still need to consider differences in analytical approaches and in cases where protein quantifications obtained by orthogonal methods differ, *cis*-pQTLs and mass spectrometry validation of probe targets may be good indicators of platform specificity^34^.

We demonstrate that proteins that are secreted, tissue-specific, more tolerant to LoF variants and with few connections in protein networks were most likely to be genetically controlled. This pattern was mainly driven by *cis* acting signals and not as apparent for the *trans* effects on protein levels, illustrating that *cis*- and *trans*-signals for serum proteins arose by different means and may differ in evolutionary properties. Our results are consistent with the notion that evolutionary important, and likely disease-relevant, genes undergo a negative selection against genetic *cis*-variants, which has been proposed as an explanation of the extreme polygenicity of complex traits^35^. The observed depletion of *cis*-variants among network hubs in our study are furthermore in line with the recently proposed omnigenic model^2^, which suggests that core disease genes are rarely affected directly by GWAS variants but rather through a multitude of smaller effects mediated through *cis*-regulation of peripheral genes in regulatory networks. Thus, while our results provide a map of *cis*-regulatory effects for 812 proteins, linking many of these to disease signals from GWAS studies, those without *cis*-effects may be even more important in the context of disease and should be studied further by other means. While hubs in the PPI network were depleted for any genetic signal, *trans* affected proteins showed higher degree of connectivity in the co-regulatory network compared to those with no detectable genetic signal. These findings demonstrate that the structure of the co-regulatory network is to some extent be driven by genetic variants affecting multiple proteins. We also note that unlike PPI networks constructed in solid tissues, the serum protein networks are composed of protein members synthesized across different tissues of the body and as such may reflect cross-tissue regulation^8^ or factors that affect the levels of circulating proteins independently of their origin.

Among proteins with genetic associations, we find that many have multiple genetic signals, both across different loci throughout the genome but also within a given locus as revealed by conditional analysis, indicating that allelic heterogeneity is common in loci regulating serum protein levels. Widespread allelic heterogeneity has been described for gene expression^36^ and complex traits in general^37^. For serum proteins, this may reflect the complex regulation and diverse origin of proteins in circulation, as these proteins may arise from almost any tissue of the body. Furthermore, *cis*-pQTLs show a roughly 40% overlap with gene expression QTLs^8,9^, suggesting that a large fraction of the genetic effect is mediated through any of the many post-transcriptional steps involved in protein maturation.

The integration of well-established genetic associations for 81 diseases and disease-related traits revealed a profound overlap with the genetic signals affecting protein levels in our study, where a third of the identified loci regulating serum protein levels colocalized with at least one GWAS phenotype. We identify examples of disease-associated loci colocalizing with many proteins, especially loci that also exhibit pleiotropic phenotype associations. Thus, it seems likely that the more complex the molecular consequences of a variant, the more likely it is to be associated with many different phenotypes, which has also been observed at the transcriptomic level^38^. The serum protein changes associated with any given disease signal can shed new light on the underlying pathways that are affected either before or after the onset of disease. The deep phenotyping of the AGES cohort allowed for an integrative analysis of genetic variants, serum protein measurements and phenotypes within the same population. For proteins regulated by loci linked to a given disease-relevant phenotype, we observed an enrichment for associations to the same phenotype measures in our cohort, thus pointing to many novel candidate proteins that may play a role in regulating or responding to these phenotypes. However, it should be noted that while a pQTL that colocalizes with a signal for a disease or clinical trait may implicate causal candidates for mediating the genetic risk, it may just as well indicate downstream events or even unrelated parallel effects of a pleiotropic variant. Furthermore, the plasma proteome has been shown to change in waves throughout the human lifespan^39^, with a large proportion of proteins changing in old age. Thus some of the associations observed in the elderly AGES cohort may not be directly transferable to a younger population, but may at the same time shed light on the physiological relevance of circulating proteins in the aging process. Our study provides genetic instruments for further studies of causal relationships for specific examples, however mechanistic and experimental studies are warranted for determining the underlying chains of events behind these complex associations. Our results offer an in-depth inventory of information regarding the interconnections between genetic variants, serum proteins and disease relevant traits, which may encourage discoveries of novel therapeutic targets and fluid biomarkers, providing a robust framework for understanding the pathobiology of complex disease.

## Methods

### The AGES cohort

Cohort participants aged 66 through 96 were included from the AGES-Reykjavik Study^40^, a prospective study of deeply phenotyped individuals of Northern European ancestry (Table S1). Blood samples were collected at the baseline visit after overnight fasting and serum lipids, glucose, HbA1c, insulin, uric acid and urea measured using standard protocols. LDL and total cholesterol levels were adjusted for statin use, with an approach similar to what has previously been described^41^. Hypertension medication use was accounted for by adding 15 mmHG to systolic blood pressure and 10 mmHG to diastolic blood pressure^42^. Serum creatinine was measured with the Roche Hitachi 912 instrument and estimated glomerular filtration rate (eGFR) derived with the four-variable MDRD Study equation^43^. Type 2 diabetes was defined from self-reported diabetes, diabetes medication use or fasting plasma glucose ≥ 7 mmol/L. Type 2 diabetes patients were excluded from all analyses for fasting glucose, fasting insulin and HbA1c. Coronary artery disease was determined using hospital records and/or cause of death registry data. A coronary artery disease event was any occurrence of myocardial infarction, ICD-10 codes: I21-I25, coronary revascularization (either CABG surgery or percutaneous coronary intervention (PCI)) or death from CHD according to a complete adjudicated registry of deaths available from the national mortality register of Iceland (ICD-10 codes I21–I25). Prostate cancer diagnosis was obtained from medical records (ICD-10 code C61). Information on migraine, Parkinson’s disease, eczema and thyroid disease was obtained from questionnaires. Alzheimer’s disease was determined with a consensus diagnosis based on international guidelines was made by a panel that includes a geriatrician, neurologist, neuropsychologist, and neuroradiologist and defined according to the criteria of the National Institute of Neurological and Communicative Disorders and Stroke and the Alzheimer’s Disease and Related Disorders Association (NINCDS-ADRDA), as previously described^44^. Hospital- and mortality data was also used to identify cases according to the ICD-10 code F00. Age-related macular degeneration (AMD) in the AGES-Reykjavik study has been previously described^45^, but in short was defined by the presence of any soft drusen and pigmentary abnormalities (increased or decreased retinal pigment) or the presence of large soft drusen ≥125μm in diameter with a large drusen area >500μm in diameter or large ≥125μm indistinct soft drusen in the absence of signs of late AMD. Maximum grip strength of the dominant hand was measured by a computerised dynamometer, as previously described^46^. Bone mineral density was estimated from a CT scan of the femur^47^. The AGES-Reykjavik study was approved by the NBC in Iceland (approval number VSN-00-063), and by the National Institute on Aging Intramural Institutional Review Board, and the Data Protection Authority in Iceland. All participants provided informed consent.

### Protein measurements

Serum levels of 4,135 human proteins, targeted by 4,782 SOMAmers^48^, were determined at SomaLogic Inc. (Boulder, US) in samples from 5,457 AGES-Reykjavik participants as previously described^8^. A few SOMAmers are annotated to more than one gene, for example when the target is a protein complex, thus the 4,782 SOMAmers are annotated to a total of 4,118 unique targets (annotated as one or more Entrez gene symbols) in the most up to date inhouse annotation database, which were used in all analyses. Sample collection and processing for protein measurements were randomized and all samples run as a single set. The SOMAmers that passed quality control had median intra-assay and inter-assay coefficient of variation (CV) <5% similar to that reported on variability in the SOMAscan assays^49^. In addition to multiple types of inferential support for SOMAmer specificity towards target proteins including cross-platform validation and detection of *cis*-acting genetic effects^8^, direct measures of the SOMAmer specificity for 779 of the SOMAmers in complex biological samples was performed using tandem mass spectrometry^8^. Previous studies have shown that pQTLs replicate well across proteomics platforms^8,9^. While a recent comparisons of protein measurements across different platforms showed a wide range of correlations^21,34^, *cis* pQTLs and validation by mass spectrometry were predictive of a strong correlation across platforms and are likely good indicators of platform specificity when protein concentrations obtained by orthogonal methods differ^34^. Hybridization controls were used to correct for systematic variability in detection and calibrator samples of three dilution sets (40%, 1% and 0.005%) were included so that the degree of fluorescence was a quantitative reflection of protein concentration. In the main text the results are described at a protein level instead of SOMAmer level, to avoid overcounting as some proteins are targeted by more than one SOMAmer that were selected to different forms or domains of the same protein. Thus, when we refer to a protein having a genetic signal, this indicates that any of the protein’s SOMAmers are associated with that genetic signal.

### Genotyping and imputation

Within the AGES cohort, 3,219 individuals were genotyped with the Illumina hu370CNV array and 2,705 individuals genotyped with the Illumina Infinium Global Screening Array. Data from both genotype arrays underwent quality control procedure, separately, removing variants with call rate < 95% and HWE p-value < 1×10^−6^. Both arrays were imputed against the Haplotype Reference Consortium imputation panel r1.1 with the Minimac3 software^50^. Post-imputation quality control consisted of filtering out variants with imputation quality R^2^ < 0.7, MAF < 0.01, as well as monomorphic and multiallelic variants for each platform separately. Genotypes for remaining variants, with matching location and alleles between platforms, were merged to create a dataset with 7,506,463 variants for 5,656 individuals (268 individuals were genotyped on both platforms, with a 99% match of genotypes for the final set of variants between platforms). The quality control procedure was performed using bcftools (v1.9)^51^ and PLINK 1.9^52^. All positions are based on genome assembly GRCh37.

### GWAS and conditional analysis

Box-Cox transformation was applied on the protein data^53^ and extreme outlier values were excluded, defined as values above the 99.5th percentile of the distribution of 99th percentile cutoffs across all proteins after scaling, resulting in the removal of an average 11 samples per SOMAmer, as previously described^18^. Within the AGES cohort, 5,368 individuals had both genetic data and protein measurements. With that sample set, 7,506,463 variants were tested for association with each of the 4,782 SOMAmers separately, in a linear regression model with age, sex, 5 genetic principal components and genotyping platform as covariates using PLINK 2.0. To obtain independent genetic signals, we performed a stepwise conditional association analysis for each SOMAmer separately with the GCTA-COJO software^54,55^. We conditioned on the current lead variant, defined as the variant with the lowest p-value, and then kept track of any new lead variants with study-wide-significant associations. Variants in strong LD (r^2^ > 0.9) with previously chosen lead variants were not considered for joint analysis to avoid multicollinearity. Associations with independent lead variants within 300kb window of the gene boundaries of the protein-coding gene were defined as *cis*-signals, and otherwise in *trans*. To compare independent signals between SOMAmers, we define any signals with lead variants in strong LD (r^2^ > 0.9) as the same signal. Due to the complex LD structure and high pleiotropy of the MHC region^56^ (chr.6, 28.47-34.45Mb) we collapsed all signals within that region to a single signal. To define loci harboring independent signals, we defined a 300 kb window around each independent signal (150 kb up- and downstream of lead variants) and collapsed all such intersecting windows. Therefore, the definition of loci is solely based on physical distances while the definition of independent signals is solely based on LD structure. The GWAS results were visualised using Circos^57^. Pathway enrichment was performed using gProfiler^58^, using the full set of measured proteins as background and considering Benjamini-Hochberg FDR<0.05 as statistically significant. Enrichment of tissue-elevated gene expression was performed using data from the Human Protein Atlas^59^ with a Fisher’s exact test, considering Benjamini-Hochberg FDR<0.05 as statistically significant.

### Comparison with previous proteogenomic studies

To evaluate the novelty of the genetic associations identified in the current study, we compared our results to 20 previously published proteogenomic studies (Supplementary Table 5), including the protein GWAS in the INTERVAL study^9^, our previously reported genetic analysis of 3,219 AGES cohort participants8, and a recent Illumina exome array analysis in 5,343 AGES participants20. In a previous proteogenomic analysis of AGES participants^8^, one *cis* variant was reported per protein using a locus-wide significance threshold, as well as *cis*-to-*trans* variants at a Bonferroni corrected significance threshold, whereas the more recent exome-array analysis^20^ reported results at a study-wide significant threshold (P<1×10^−10^). Due to these differences in reporting criteria, we only considered the associations in previous AGES results that met the current study-wide p-value threshold (P < 1.046×10^−11^). For all other studies we retained the pQTLs at the reported significance threshold. In addition, we performed a lookup of all independent pQTLs from the current study available in summary statistics from the INTERVAL study, considering them known if they reached a study-wide significance in their data. We calculated the LD structure between the reported significant variants for all studies, using 1000 Genomes v3 EUR samples, but using AGES data when comparing to previously reported AGES results. We considered variants in LD (r^2^>0.9 for consistency for defining signals across SOMAmers described above, but results for r^2^>0.5 are additionally shown in Supplementary Note 1) to represent the same signal across studies. Comparison was performed on protein level, by matching the reported Entrez gene symbol from each study.

### Enrichment analysis

We grouped the proteins into three categories derived from our GWAS results; a) proteins with at least one *cis* signal, b) proteins with no *cis* signals and at least one *trans* signal and c) proteins with no genetic signal. From our data we also derived three continuous traits for a given protein; a) number of associated independent signals, b) highest absolute beta coefficient of all associated signals and c) number of proteins that share genetic signals with the given protein, which is essentially a quantitative representation of whether a protein is a part of a *trans* hotspot. We fetched publicly available data regarding; a) tissue elevated gene expression, where “Tissue Enriched” in our analyses refers to the “Tissue Enriched”, “Tissue Enhanced” or “Group Enriched” categories defined by Uhlen et al.^22^, b) tissue elevated protein expression, where “Tissue Enriched” in our analyses refers to the “Tissue Enriched”, “Tissue Enhanced” or “Group Enriched” categories defined by Wang et al.^23^, c) annotation of secreted and transmembrane proteins, classifying proteins as secreted or transmembrane if it was predicted so by at least one method or one segment, respectfully^22^, d) gene-level loss-of-function intolerance^24^ and e) network degree in the InWeb protein-protein interaction network^25^. Furthermore, we estimated hub status of proteins within the serum protein co-regulation network derived from the AGES cohort^8^. Protein classifications were compared using a Fisher’s exact test, where the estimate is the odds ratio. Continuous parameters were compared between protein classes using the Wilcoxon Rank Sum test and for the estimate we calculated the median of the difference between values from the two classes, so the size of the estimate is dependent on the scale of the values. For comparing two continuous traits we used Spearman’s Rho correlation. We report 95% confidence intervals of all estimates.

### GWAS colocalization analysis

We included 81 phenotypic traits including major disease classes in the colocalization analysis, for which GWAS summary statistics were publicly available from consortium websites and the GWAS catalog^60^. We restricted the study selection to those with study sample sizes of n > 10K, of primarily European Ancestry (to match the AGES cohort’s LD structure), having at least one genome-wide significant association (P<5×10^−8^) and selecting one study per phenotype (Table S8). For each trait, significant loci were defined by identifying all genome-wide variants (P<5×10^−8^) at least 500kb apart, defining a flanking region of 1 Mb around each lead variant and finally merging overlapping regions. For each GWAS locus, all SOMAmers with a study-wide significant association (*cis* or *trans*) within the given region were tested for colocalization, if at least 50 SNPs in the region had complete information from both trait and protein GWAS. When the MAF was not available for a given GWAS, the 1000 Genomes EUR MAF was used instead. Colocalization analysis was performed with coloc (v.3.2-1)^61^, using the coloc.abf function with default priors. High and medium colocalization support was defined as PP.H4>0.8 and PP.H4>0.5, respectively. Conditional colocalization analysis was performed using coloc 4.0-4^62^, using the “allbutone” option and restricted to loci harboring more than one independent signal per protein. Unlike the primary coloc analysis, the conditional analysis requires the GWAS effect size to be included, thus the phenotypes AMD, ATD and PD were excluded from this analysis which did not have this information available in the GWAS summary statistics. Results were visualized with LocusCompare^63^.

### Phenotype associations

For each GWAS phenotype with a corresponding measurement in AGES and well represented at the population level (Table S8), the colocalized proteins were tested for association with the phenotype in all AGES participants with protein data available (n = 5,457, see n missing per phenotype in Table S1), in a linear or logistic regression model adjusted for age and sex. The SOMAmer with the lowest P-value was chosen for each protein, and P-values were subsequently adjusted for the number of proteins tested for each trait by Benjamini-Hochberg FDR. For each phenotype with at least five colocalized proteins, the proportion of significantly associated proteins (FDR<0.05) was compared to that obtained by 1000 randomly sampled protein sets of the same size, again choosing the SOMAmer with the lowest P-value per protein, and an empirical P-value calculated. The analysis was repeated by excluding proteins originating from loci where five or more proteins colocalized with the same phenotype. The same enrichment analysis was additionallly performed for each individual locus where where five or more proteins colocalized with the same phenotype.

## Acknowledgements

The authors thank the staff of the Icelandic Heart Association for their contribution to AGES-Reykjavik and all study participants for their invaluable contributions to this study.

The study was funded by Icelandic Heart Association contract HHSN271201200022C, National Institute on Aging contract N01-AG-12100, and Althingi (the Icelandic Parliament). V.E. and Va.G. are supported by the Icelandic Research Fund (IRF grants 195761-051, 184845-053 and 206692-051) and Va.G. holds a postdoctoral research grant from the University of Iceland Research Fund.

## Author Contributions

A.G., Va.G., V.E., and Vi.G designed the study. A.G., Va.G., G.T.A., E.F.G., B.G.J. and T.A. performed data analysis. J.R.L. and L.L.J. provided expertise on proteomics data and contributed to discussion. Vi.G. and V.E. supervised the project. A.G. and Va.G. wrote the first draft of the manuscript, with all coauthors contributing to data interpretation, manuscript editing, and revision.

## Declaration of Interests

The study was supported by the Novartis Institute for Biomedical Research, and protein measurements for the AGES-Reykjavik cohort were performed at SomaLogic. J.R.L. and L.L.J. are employees and stockholders of Novartis. All other authors have no conflict of interests to declare.

## Data Availability

The custom-design Novartis SOMAscan is available through a collaboration agreement with the Novartis Institutes for BioMedical Research. Data from the AGES Reykjavik study are available through collaboration under a data usage agreement with the IHA. All other data supporting the conclusions of the paper are presented in the main text and supplementary materials.

