## Supplementary Material for "A genome-wide association study of serum proteins reveals shared loci with common diseases"

### The Serum Proteome Links Genetics to Complex Disease

[1]Icelandic Heart Association, Holtasmari 1, 201 Kopavogur, Iceland.
[2]Faculty of Medicine, University of Iceland, 101 Reykjavik, Iceland.
[3]Laboratory of Epidemiology and Population Sciences, Intramural Research Program, National Institute on Aging, Bethesda, MD 20892-9205, USA.
[4]GNF Novartis, 10675 John Jay Hopkins Drive, San Diego, CA 92121, USA.
[5]Novartis Institutes for Biomedical Research, 22 Windsor Street, Cambridge, MA 02139, USA.

*These authors contributed equally as joint-first authors
[#]These authors contributed equally as joint-senior authors

### Supplementary Note 1: Novelty estimates for pQTLs reported in the current study

The current study is a genome-wide analysis of 7.5M variants imputed using the HRC reference panel in 5,368 AGES participants and measurements of 4,782 SOMAmers. We compared the pQTLs identified in the current study to previously reported studies: a) the INTERVAL-study GWAS[1] of 3,283 SOMAmers in 3,301 individuals based on 10.6 million imputed (combined 1000 Genomes Phase 3-UK10K reference panel) autosomal variants, b) our previously published[2] *cis* pQTL (300kb window up- and downstream of gene boundaries) and *cis*-to-*trans* pQTLs analysis for 4,783 SOMAmers in 3,200 AGES participants using genetic variants imputed using the 1000 Genomes v3 reference panel and c) a recent Illumina exome array analysis[3] of 54,469 variants and 4,782 SOMAmers in up to 5,343 AGES participants, d) 17 additional proteogenomic studies with a more limited protein coverage (Supplementary Table 5).

In the current study, genetic signals were defined as shared across SOMAmers if the lead variants were in strong LD ($r^2$>0.9). Therefore, the same LD threshold was used to define shared genetic signals across studies based on reported lead variants. However, the study comparison was also performed using a more stringent threshold ($r^2$>0.5). In addition, we performed a lookup of lead independent variants in the current study in the publicly available summary statistics from the INTERVAL study[1].

As shown in Supplementary Note Table 1 and Supplementary Fig. 4, 1,527 (37.1%) associations were considered novel at the primary LD threshold and 1,261 (30.7%) remained so at the more stringent LD threshold. The majority of proteins (1,927 or 91.8%) for which we find a pQTL have a previously reported pQTL, with a large proportion originating from our recent exome chip study (Emilsson et al. 2020), while the current study increases considerably the number of known genetic signals for serum protein levels (39.3% of genetic signals defined as novel at the primary LD threshold).

Finally, we considered novelty at the locus level, in addition to an evaluation based on the independent signals. Combining independent signals within 300kb of each other, we have 799 loci associated with protein levels. Of those, 107 (13.3%) are novel, in the sense that they are >500kb away from lead pQTL variants reported in other studies. Of 3,131 locus-protein associations, 607 (19.4%) are previously unreported. If we exclude our own exome array analysis in the AGES cohort[3], we find that 18.1% of the loci are novel and 40.8% of the locus-protein associations.

**Supplementary Note Table 1 –** Comparison of independent pQTLs (association, genetic signal and protein) identified in the current study compared to previously reported proteogenomic studies. The comparison is shown at two different LD thresholds.

| LD threshold (r2) | Study compared to | Type | Total | Known | | Addition | | Novel | |
|---|---|---|---|---|---|---|---|---|---|
| | | | count | count | % | count | % | count | % |
| 0.9 | Sun et al. 2018 | assoc | 4113 | 1305 | 31.7 | NA | NA | 2808 | 68.3 |
| 0.9 | Sun et al. 2018 | signal | 2087 | 707 | 33.9 | 189 | 9.1 | 1191 | 57.1 |
| 0.9 | Sun et al. 2018 | protein | 2099 | 387 | 18.4 | 688 | 32.8 | 1024 | 48.8 |
| 0.9 | Emilsson et al. 2018 | assoc | 4113 | 1087 | 26.4 | NA | NA | 3026 | 73.6 |
| 0.9 | Emilsson et al. 2018 | signal | 2087 | 428 | 20.5 | 55 | 2.6 | 1604 | 76.9 |
| 0.9 | Emilsson et al. 2018 | protein | 2099 | 485 | 23.1 | 766 | 36.5 | 848 | 40.4 |
| 0.9 | Emilsson et al. 2020 | assoc | 4113 | 1573 | 38.2 | NA | NA | 2540 | 61.8 |
| 0.9 | Emilsson et al. 2020 | signal | 2087 | 355 | 17.0 | 30 | 1.4 | 1702 | 81.6 |
| 0.9 | Emilsson et al. 2020 | protein | 2099 | 729 | 34.7 | 998 | 47.5 | 372 | 17.7 |
| 0.9 | Other | assoc | 4113 | 333 | 8.1 | NA | NA | 3780 | 91.9 |
| 0.9 | Other | signal | 2087 | 202 | 9.7 | 232 | 11.1 | 1653 | 79.2 |
| 0.9 | Other | protein | 2099 | 73 | 3.5 | 396 | 18.9 | 1630 | 77.7 |
| 0.9 | Any | assoc | 4113 | 2586 | 62.9 | NA | NA | 1527 | 37.1 |
| 0.9 | Any | signal | 2087 | 1060 | 50.8 | 206 | 9.9 | 821 | 39.3 |
| 0.9 | Any | protein | 2099 | 1016 | 48.4 | 911 | 43.4 | 172 | 8.2 |
| 0.5 | Sun et al. 2018 | assoc | 4113 | 1346 | 32.7 | NA | NA | 2767 | 67.3 |
| 0.5 | Sun et al. 2018 | signal | 2087 | 734 | 35.2 | 246 | 11.8 | 1107 | 53.0 |
| 0.5 | Sun et al. 2018 | protein | 2099 | 405 | 19.3 | 670 | 31.9 | 1024 | 48.8 |
| 0.5 | Emilsson et al. 2018 | assoc | 4113 | 1252 | 30.4 | NA | NA | 2861 | 69.6 |
| 0.5 | Emilsson et al. 2018 | signal | 2087 | 522 | 25.0 | 92 | 4.4 | 1473 | 70.6 |
| 0.5 | Emilsson et al. 2018 | protein | 2099 | 531 | 25.3 | 720 | 34.3 | 848 | 40.4 |
| 0.5 | Emilsson et al. 2020 | assoc | 4113 | 2010 | 48.9 | NA | NA | 2103 | 51.1 |
| 0.5 | Emilsson et al. 2020 | signal | 2087 | 636 | 30.5 | 58 | 2.8 | 1393 | 66.7 |
| 0.5 | Emilsson et al. 2020 | protein | 2099 | 920 | 43.8 | 807 | 38.4 | 372 | 17.7 |
| 0.5 | Other | assoc | 4113 | 438 | 10.6 | NA | NA | 3675 | 89.4 |
| 0.5 | Other | signal | 2087 | 267 | 12.8 | 300 | 14.4 | 1520 | 72.8 |
| 0.5 | Other | protein | 2099 | 104 | 5.0 | 365 | 17.4 | 1630 | 77.7 |
| 0.5 | Any | assoc | 4113 | 2852 | 69.3 | NA | NA | 1261 | 30.7 |
| 0.5 | Any | signal | 2087 | 1234 | 59.1 | 203 | 9.7 | 650 | 31.1 |
| 0.5 | Any | protein | 2099 | 1161 | 55.3 | 766 | 36.5 | 172 | 8.2 |

### Supplementary Note 2: Replication of pQTLs between INTERVAL and AGES cohorts

*Replication of INTERVAL study pQTLs in AGES*

We performed a lookup of previously reported pQTLs from the INTERVAL study[1] (n = 3,301). The comparison was based on the reported independent variants from a conditional analysis, however comparing univariate effect sizes and P-values to our GWAS results for consistency.

Replication of pQTLs from the INTERVAL study was good, with 84.6% of reported associations directionally consistent in AGES and 75.6% both directionally consistent and nominally significant (Supplementary Note Table 2, Supplementary Fig. 5). While the proportion of study-wide significant associations in AGES increased with a stronger reported significance in the INTERVAL study as expected, rather surprisingly the proportion of non-significant associations remained constant around 20% independent of the reported significance (Supplementary Fig. 5C). This was explained by a large proportion of pQTLs reported in the INTERVAL study that are located on chromosome 19, where a strikingly high proportion was not replicated in AGES (Supplementary Fig. 6A). A further investigation narrowed this region down to the *NLRP12* locus (chr 19, 54,319,624-54,327,869), a *trans* hotspot reported in the INTERVAL study with a total of 391 associations, whereof only 20 were nominally significant in the current study (Supplementary Fig. 4B), with the minimum observed P-value = 0.002 in AGES at this locus compared to a minimum P-value = $6.5\times10^{-244}$ in the INTERVAL study. Given the strong observed significance in the INTERVAL study, the associations at this locus are unlikely to be false positives, however the discrepancies between the two studies at this locus may rather reflect some cohort-specific attributes, as for instance the INTERVAL study is based on plasma samples compared to serum samples in AGES and there is a large age gap between the two cohorts. More specifically, the INTERVAL cohort consists of young (mean age ~44 years), healthy blood donors of European descent and recruited in England and the proteins are measured in plasma samples, while the population-based AGES cohort consists of elderly (mean age ~76 years) Icelanders, many of whom suffer from chronic diseases and the proteins are measured in serum samples. Others have suggested that differences in observed pQTLs in this locus may be driven by sample handling and white blood cell lysis[4]. The lead variant in the INTERVAL study at this locus (rs62143197) is an intron variant in *NLRP12*, which is also an eQTL for *NLRP12* in GTEx (P = $3.2\times10^{-15}$ for whole blood, https://gtexportal.org/home/). A lookup of this variant in UKBB associations[5] (http://geneatlas.roslin.ed.ac.uk/) revealed strong associations with monocyte percentage (beta = -0.12, P = $3.4 \times 10^{-155}$) and counts (beta = -0.009, P = $1.5 \times 10^{-141}$).

When the *NLRP12* locus was excluded from the pQTL comparison, 97.6% of associations reported in the INTERVAL study were directionally consistent in the current study and 93.3% were both directionally consistent and nominally significant (Supplementary Note Table 2, Supplementary Fig. 5).

*Replication of AGES pQTLs in the INTERVAL study*

We next performed a lookup of the pQTLs identified in the current study (n = 5,368) in summary statistics from the INTERVAL study[1] (n = 3,301). The comparison was based on the lead independent variants from the conditional analysis, however comparing univariate effect sizes and P-values for consistency between the two studies and restricted to the 4,082 associations that were study-wide significant in both the joint model (from the conditional analysis) and the univariate GWAS in AGES.

Of the 2,716 associations that could be compared between the studies, we found 94.1% to be directionally consistent and 81.2% were both directionally consistent and at least nominally significant ($P < 0.05$) in the INTERVAL study (Supplementary Note Table 2). When we restricted the comparison to 668 pQTLs defined as novel in the current study (see Supplementary Note 1) and with information available in the INTERVAL study, we still found 89.8% to be directionally consistent and 62.6% to be both directionally consistent and nominally significant ($P < 0.05$).

**Supplementary Note Table 2** – Overview of reported independent pQTLs in the INTERVAL study (Sun et al. 2018) with their replication status in AGES and vice versa. The INTERVAL study results are also shown excluding the *NLRP12* locus, a *trans* hotspot in the INTERVAL study. For the current AGES study, we show the results for all associations and only those defined as novel.

| Discovery<br>Replication | Sun et al. 2018<br>AGES | Sun et al. 2018 (excluding the NLRP12 locus)<br>AGES | AGES<br>Sun et al. 2018 | AGES (novel only)<br>Sun et al. 2018 |
|---|---|---|---|---|
| Associations (n) | 1905 | 1514 | 2716 | 668 |
| SNPs (n) | 1046 | 1037 | 1576 | 487 |
| SOMAmers (n) | 1359 | 1022 | 1611 | 543 |
| Directionally consistent (n (%)) | 1611 (84.6) | 1476 (97.5) | 2555 (94.1) | 600 (89.8) |
| Nominally significant (P<0.05) in replication (n (%)) | 1441 (75.6) | 1421 (93.9) | 2227 (82.0) | 420 (62.9) |
| Directionally consistent and nominally significant (P<0.05) in replication (n (%)) | 1417 (74.4) | 1413 (93.3) | 2206 (81.2) | 418 (62.6) |

### Supplementary Note 3: GWAS of the serum protein co-regulatory network

We regressed 54 Eigenproteins (1st and 2nd PCs) on 7.5 million variants that were assayed or imputed in all 5,368 AGES participants with protein data. Combined, the 1st and 2nd PCs capture on average 45% of the variance in each serum protein module. We assumed an additive linear model for each module's Eigenprotein. Applying the conventional P-threshold of $5.0\times10^{-8}$ for genome-wide significance when a linkage disequilibrium (LD) $r^2<0.8$ is used for independent variants, we find that 24 (89%) out of 27 modules are associated with at least one independent network-associated protein SNP (npSNP) (Supplementary Note Table 3). This is a marked increase (26%) in number of identified npSNPs compared to previous results based on 1.5 million markers in 3,219 AGES participants[2]. Applying a naive conservative Bonferroni correction for number of variants and Eigenproteins tested, 18 (67%) modules showed significant association to at least one independent npSNP at $P<1.2\times10^{-10}$. Often, certain npSNPs are associated with more than one protein module that is consistent with their relationship[2].

When the significant npSNPs listed in Supplementary Note Table 3 were tested against all proteins we considered associations at $P < 1\times10^{-7}$ significant, i.e. using Bonferroni correction for number of npSNPs and proteins tested. The majority of npSNPs affected many individual proteins in *cis* and/or *trans* (mostly *trans*), while the proteins that were affected comprised the module(s) associated with the corresponding npSNP which is in line with previous observations[2]. For instance, four npSNPs across three chromosomal sites including a region harboring *APOE/TOMM40*, were associated with either 1$^{st}$ or 2$^{nd}$ PCs for the protein module 11 (PM11) (Supplementary Note Table 3). These variants regulated 136 proteins in *cis* and/or *trans* including 96% of all proteins that constitute PM11. These results show that npSNP affect many co-regulated proteins and are underlying the architecture of the serum protein network.

The npSNPs associated with the module PM11, are also known variants in genome-wide association studies (GWAS) linked to Late-Onset Alzheimer´s Disease (LOAD), lipoproteins and coronary heart disease (CHD) (Supplementary Note Table 3). In fact, the majority of the npSNPs listed in Supplementary Note Table 3 have previously been linked to an array of clinical traits and complex diseases. For instance, a recent GWAS study identified a number of variants associated with immunoglobulin G N-glycosylation (IgGG)[6], but aberrant glycosylation of IgG has been linked to a number of age-related disease outcomes[7]. Here we find that the npSNPs rs35590487 and rs35592422 associated with the protein module PM20 (Supplementary Note Table 3), are also found to be associated with IgGG at chromosome 14[6,8]. More to the point, rs35590487 and rs35592422 regulate 37 proteins including 56% of all proteins that comprise PM20. Other striking examples include the npSNPs associated with the 1$^{st}$ or 2$^{nd}$ PCs of the protein modules PM1, PM7, PM12-PM15 (Supplementary Note Table 3), but these are known GWAS risk variants for age-related macular degeneration (AMD)[9]. These and other examples[2], highlight the genetic architecture of the serum protein network and demonstrate that the protein network and the genetic risk of diseases is intimately connected.

**Supplementary Note Table 3. Genetic variants associated with module $E^{(q)}$s**

Identification of genetic variants associated with different modules $E^{(q)}$s (q is the annotation for a specific module). Associations at the conventional genome-wide significant $P<5.0\times10^{-8}$ or naïve Bonferroni corrected $P<1.2\times10^{-10}$ are reported. N/A, not applicable. GWAS, phenotypes associated with the corresponding npSNP at $P<5.0\times10^{-8}$ as reported in the PhenoScanner and/or the GWAS catalogue.

| Module | Cluster | 1st $E^{(q)}$ npSNP | 2nd $E^{(q)}$ npSNP | npSNP chr. | P-value npSNP | N proteins affected | Known GWAS** | *Cis* expression QTL (tissue)* |
|---|---|---|---|---|---|---|---|---|
| **PM1** | I | rs537179722 | | 6 | 6.9E-12 | 36 | RA, AMD, LC | SKIV2L (many) |
| | | | rs887829 | 2 | 7.9E-43 | 14 | Bilirubin | UGT1A3 (liver) |
| **PM2** | II | rs704 | | 17 | 2.4E-10 | 846 | OPG | TMEM199 (many) |
| | | rs12439785 | | 15 | 3.2E-09 | 7 | Myopia | None |
| | | | rs704 | 17 | 1.8E-12 | 846 | OPG | TMEM199 (many) |
| **PM3** | II | rs11574452 | | 4 | 4.5E-16 | 289 | Albumin | PF4 (fibroblasts) |
| | | | rs115008613 | 1 | 9.9E-09 | None | None | None |
| | | | rs73225360 | X | 4.2E-08 | None | None | None |
| **PM4** | II | rs11574452 | | 4 | 6.7E-10 | 289 | Albumin | PF4 (fibroblasts) |
| | | | rs10761741 | 10 | 4.2E-08 | 19 | MPV, TG | NRBF2 (fibroblasts) |
| | | | rs704 | 17 | 1.6E-37 | 846 | OPG | TMEM199 (many) |
| **PM5** | II | rs2731674 | | 5 | 1.5E-09 | 115 | Height | F12 (liver) |
| | | | rs2345872 | 2 | 4.2E-08 | 8 | None | FMNL2 (fibroblasts) |
| | | | rs75931041 | 11 | 2.0E-08 | None | None | None |
| **PM6** | II | rs73088258 | | 7 | 3.8E-08 | 6 | None | None |
| **PM7** | II | rs241779 | | 17 | 1E-302 | 729 | AMD | TMEM199 (many) |
| | | | rs241775 | 17 | 1E-302 | 728 | AMD | TMEM199 (many) |
| **PM8** | II | | rs704 | 17 | 2.6E-19 | 846 | OPG | TMEM199 (many) |
| **PM9** | II | rs1178713 | | 3 | 4.1E-08 | 14 | None | None |
| **PM10** | II | rs704 | | 17 | 3.1E-72 | 846 | OPG | TMEM199 (many) |
| | | | rs704 | 17 | 7.9E-143 | 846 | OPG | TMEM199 (many) |
| | | | rs11574452 | 4 | 1.9E-11 | 289 | Albumin | PF4 (fibroblasts) |
| | | | rs597808 | 12 | 1.1E-08 | 15 | CRC, HT-H, SLE | ALDH2 (esophagus) |
| **PM11** | III | rs483082 | | 19 | 4.8E-160 | 59 | LOAD, TG | APOE (skin) |
| | | rs1355537 | | 3 | 3.1E-15 | 41 | None | BCHE (nerve) |
| | | | rs429358 | 19 | 3.0E-231 | 63 | LOAD, LDL, CHD | TOMM40 (brain) |
| | | | rs1260326 | 2 | 1.4E-09 | 26 | TG, CRP, CHOL | NRBP1 (many) |
| **PM12** | III | | rs704 | 17 | 1.9E-16 | 846 | OPG | TMEM199 (many) |
| | | | rs3117116 | 6 | 9.9E-12 | 25 | RA, MS | HLA-DRB5 (many) |
| | | | rs528298 | 1 | 5.9-09 | 198 | AMD | CFHR1/3 (many) |
| **PM13** | III | rs528298 | | 1 | 2.2E-14 | 198 | AMD | CFHR1/3 (many) |
| | | rs1042663 | | 6 | 4.5E-13 | 152 | RA, AMD, NV, MPV | SKIV2L (many) |
| | | rs171360 | | 17 | 7.8E-10 | 526 | None | TMEM97 (thyroid) |

| | | | | | | | | |
|---|---|---|---|---|---|---|---|---|
| | | | rs528298 | 1 | 5.1E-10 | 198 | AMD | CFHR1/3 (many) |
| | | | rs115009784 | 6 | 7.9E-10 | 53 | AMD, RA | DXO (many) |
| | | | rs11574452 | 4 | 2.9E-09 | 289 | Albumin | PF4 (fibroblasts) |
| **PM14** | III | rs74480769 | | 5 | 4.1E-11 | 111 | None | C7 (many) |
| | | rs1042663 | | 6 | 1.2E-10 | 152 | RA, AMD, NV, MPV | SKIV2L (many) |
| | | rs528298 | | 1 | 1.1E-09 | 198 | AMD | CFHR1/3 (many) |
| | | rs12067507 | | 1 | 2.7E-08 | 56 | None | None |
| **PM15** | III | rs1042663 | | 6 | 2.3E-17 | 152 | RA, AMD, NV, MPV | SKIV2L (many) |
| | | rs528298 | | 1 | 2.7E-17 | 198 | AMD | CFHR1/3 (many) |
| | | rs74480769 | | 5 | 2.4E-14 | 111 | None | C7 (many) |
| | | rs171360 | | 17 | 1.1E-08 | 526 | None | TMEM97 (thyroid) |
| | | | rs115008613 | 1 | 3.7E-08 | None | None | None |
| **PM16** | IV | | rs11574452 | 4 | 1.9E-09 | 289 | Albumin | PF4 (fibroblasts) |
| | | | rs61759306 | 1 | 3.9E-08 | None | RBC, MCHC | SMIM1 (blood) |
| **PM17** | IV | None | None | N/A | N/A | N/A | N/A | N/A |
| **PM18** | IV | rs35120348 | | 12 | 3.7E-09 | 5 | None | METTL21B (blood) |
| **PM19** | IV | None | None | N/A | N/A | N/A | N/A | N/A |
| **PM20** | V | rs35590487 | | 14 | 6.7E-18 | 35 | IgGG | TMEM121 (many) |
| | | rs111648045 | | 2 | 3.7E-08 | 6 (complex) | None | None |
| | | | rs35592422 | 14 | 1.7E-25 | 36 | IgGG | TMEM121 (many) |
| | | | rs2855812 | 6 | 6.0E-12 | 10 | T1D, Asthma, RA | MICB (many) |
| | | | rs55800253 | 11 | 2.8E-08 | None | None | None |
| **PM21** | V | | rs9268833 | 6 | 4.0E-09 | 13 | RA, Asthma | HLA (many) |
| **PM22** | V | | rs3756074 | 4 | 9.2E-09 | 285 | Albumin | PF4 (fibroblasts) |
| **PM23** | V | rs77869924 | | 16 | 2.8E-08 | 3 | None | None |
| | | | rs77625270 | 19 | 3.4E-23 | 1 | None | TIMM44 (blood) |
| | | | rs9411378 | 9 | 4.5E-09 | 65 | LDL, MI, VTE | ABO (many) |
| | | | rs16850360 | 4 | 4.0E-08 | 244 | CAC | PF4 (fibroblasts) |
| **PM24** | V | | rs1250258 | 2 | 1.1E-57 | 7 | CHD, Endometriosis | ATIC (blood) |
| | | | rs5985 | 6 | 5.7E-11 | 3 | ESC | None |
| | | | rs353634 | 11 | 7.8E-09 | 1 | Hair colour | CD44 (artery) |
| **PM25** | V | rs185289117 | | 14 | 3.0E-85 | 42 | None | None |
| | | rs150787591 | | 2 | 7.4E-09 | None | None | None |
| | | rs4016738 | | 3 | 3.2E-08 | 2 | MC | WDR48 (blood) |
| | | | rs80324969 | 14 | 2.5E-25 | 38 | None | CRIP2 (heart) |
| | | | rs3134998 | 6 | 2.7E-09 | 11 | RA, Asthma | HLA (many) |
| **PM26** | V | rs76833488 | | 15 | 5.7E-09 | 15 | None | None |
| | | | rs11574452 | 4 | 1.7E-13 | 289 | Albumin | PF4 (fibroblasts) |
| | | | rs4131289 | 5 | 2.1E-09 | 1 | APTT, CKD | RGS14 (many) |
| **PM27** | V | None | None | | N/A | | N/A | N/A |

*An example of *cis* expression QTL from the GTEx database.

**For a qualified proxy the correlation between npSNP and corresponding lead GWAS SNP was $r^2 \geq 0.8$. Abbreviations: RA, rheumatoid arthritis; AMD, age-related macular degeneration; LC, lymphocyte count; OPG, Osteoprotegerin levels; MPV, mean platelet volume; TG, triglyceride; CRC, colorectal cancer; HT-H, hypothyroidism; SLE, systemic lupus erythrematosus; LOAD, late-onset Alzheimer´s disease; CRP, C-reactive protein; CHOL, total cholesterol; MS, multiple sclerosis; WM, brain white matter; RBC, red blood count; MCHC, mean corpuscular hemoglobin concentration; IgGG, IgG glycosylation; T1D, type 1 diabetes; VTE, venous thromboembolism; CAC, coronary artery calcium; CHD, coronary heart disease; ESC, end-stage coagulation; MC, monocyte count; APTT, activated partial thromboplastin time ; CKD, chronic kidney disease

### Supplementary Figures

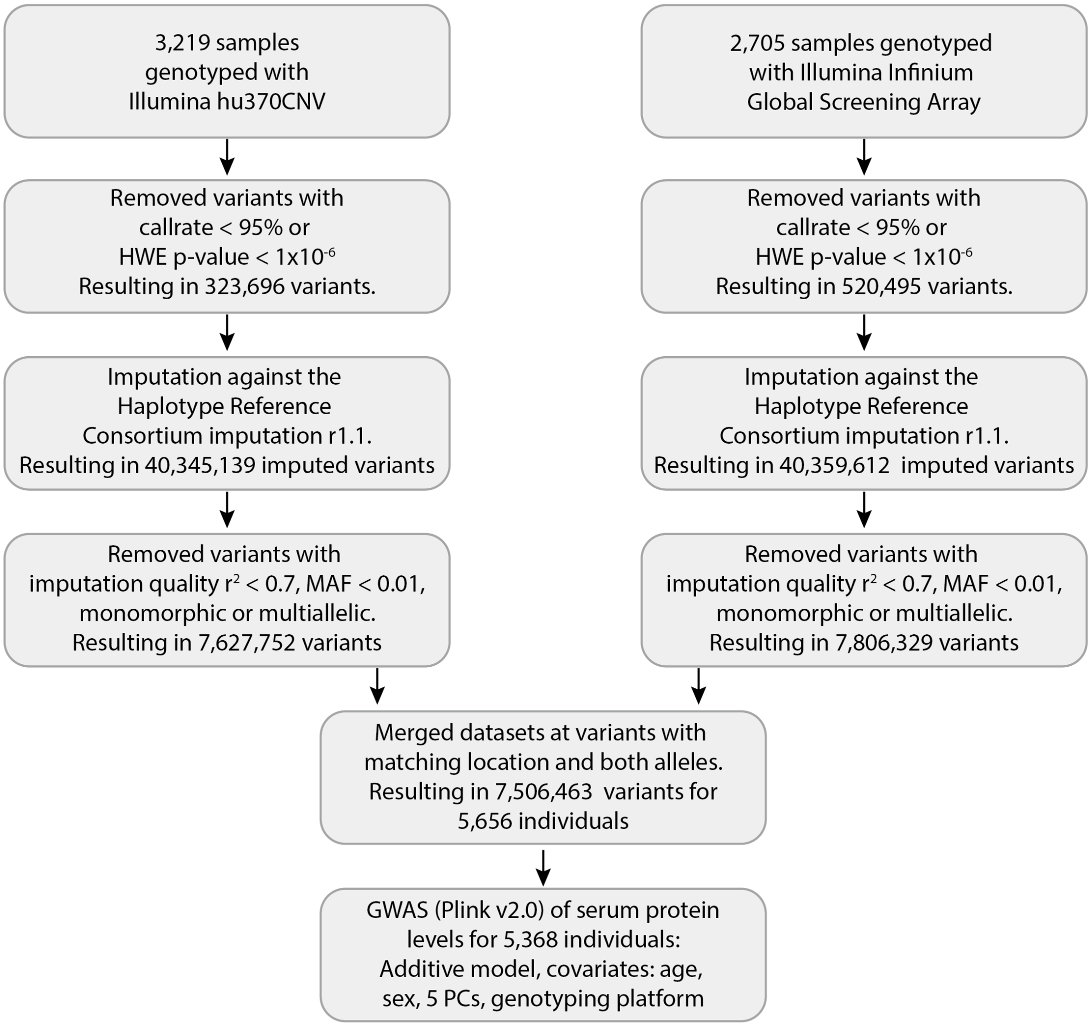

**Supplementary Figure 1** A flowchart illustrating the preprocessing steps of AGES-Reykjavik genotype data undertaken before the GWAS of serum protein levels. In short, the AGES-Reykjavik samples were genotyped using two different platforms and the resulting genotypes filtered in a similar manner before HRC imputation. The two datasets were merged after post-imputation filtering and used for the GWAS analysis, while including genotype platform among the covariates.

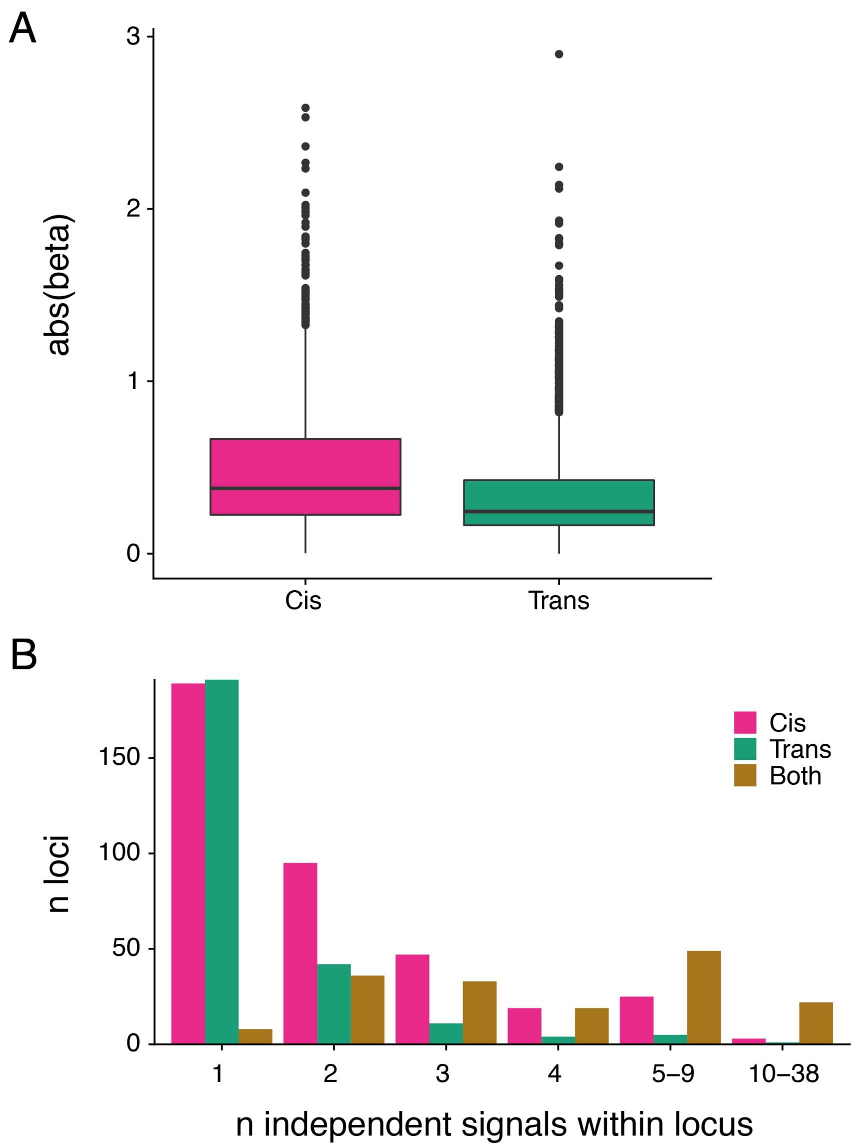

**Supplementary Figure 2** A) Boxplot comparing effect sizes (absolute beta value) between *cis* and *trans* associations. Only independent associations (Table S3) are included in the comparison and the strongest association (lowest P-value) chosen per protein if targeted by more than one SOMAmer. The median values are 0.38 (*cis*) and 0.24 (*trans*), and are significantly different (two-sided Wilcoxon's P = $6.2 \times 10^{-7}$). Boxplots indicate median value, 25th and 75th percentile, whiskers extend to smallest/largest value no further than 1.5×IQR, outliers are shown. B) Barplot showing the distribution of study-wide independent signals per locus. Independent signals within 300 kb from each other were considered to be in the same locus (see Methods for details).

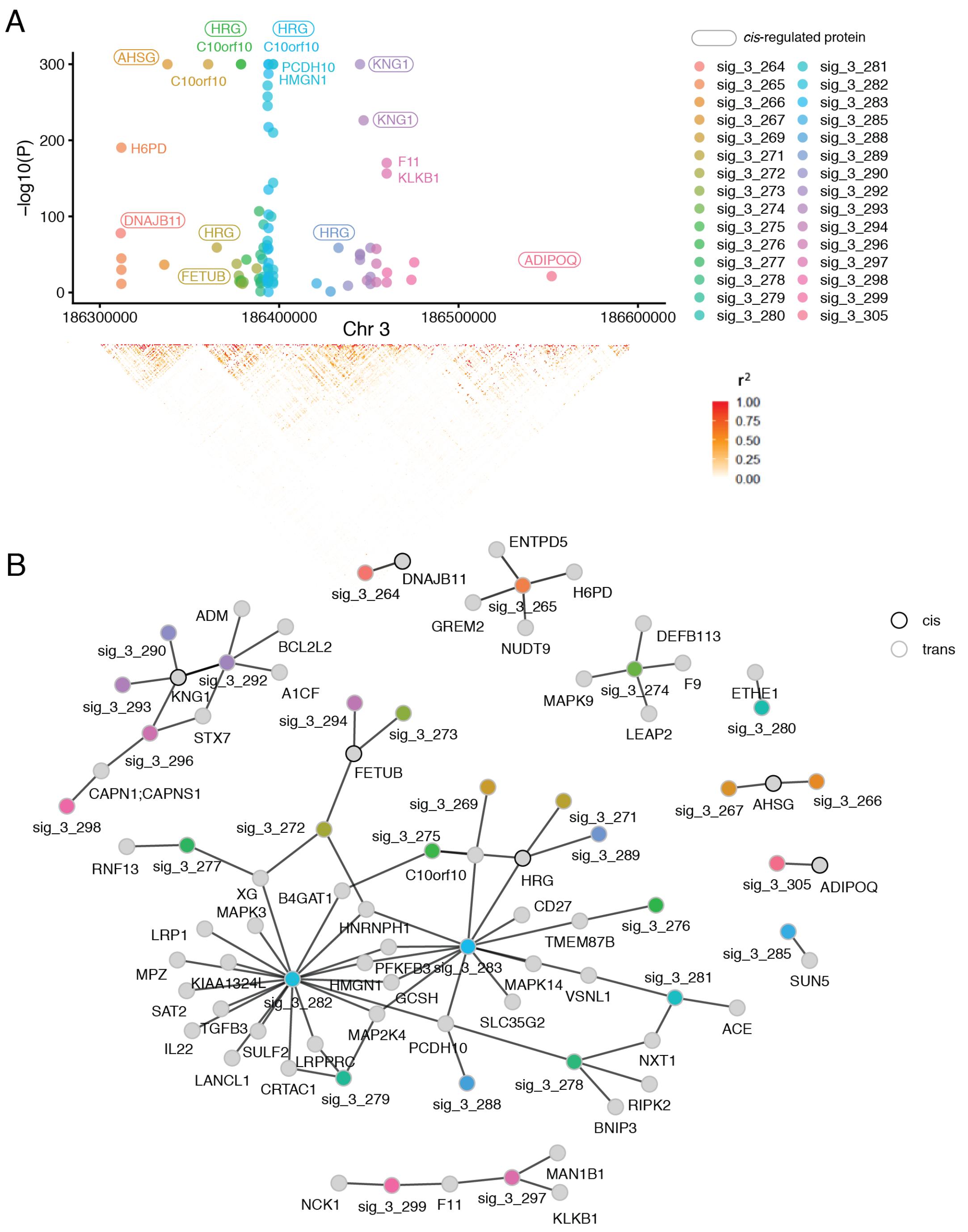

**Supplementary Figure 3** An overview of a region on chromosome 3 (186.3-186.6 Mb) where 30 independent study-wide significant signals were identified (n = 5,368). A) Each point represents the -log10(P-value) for the associations between a given independent signal and a serum protein. The LD structure in the region (based on AGES data) is shown below. B) The network demonstrates the links between independent signals in the region and serum proteins.

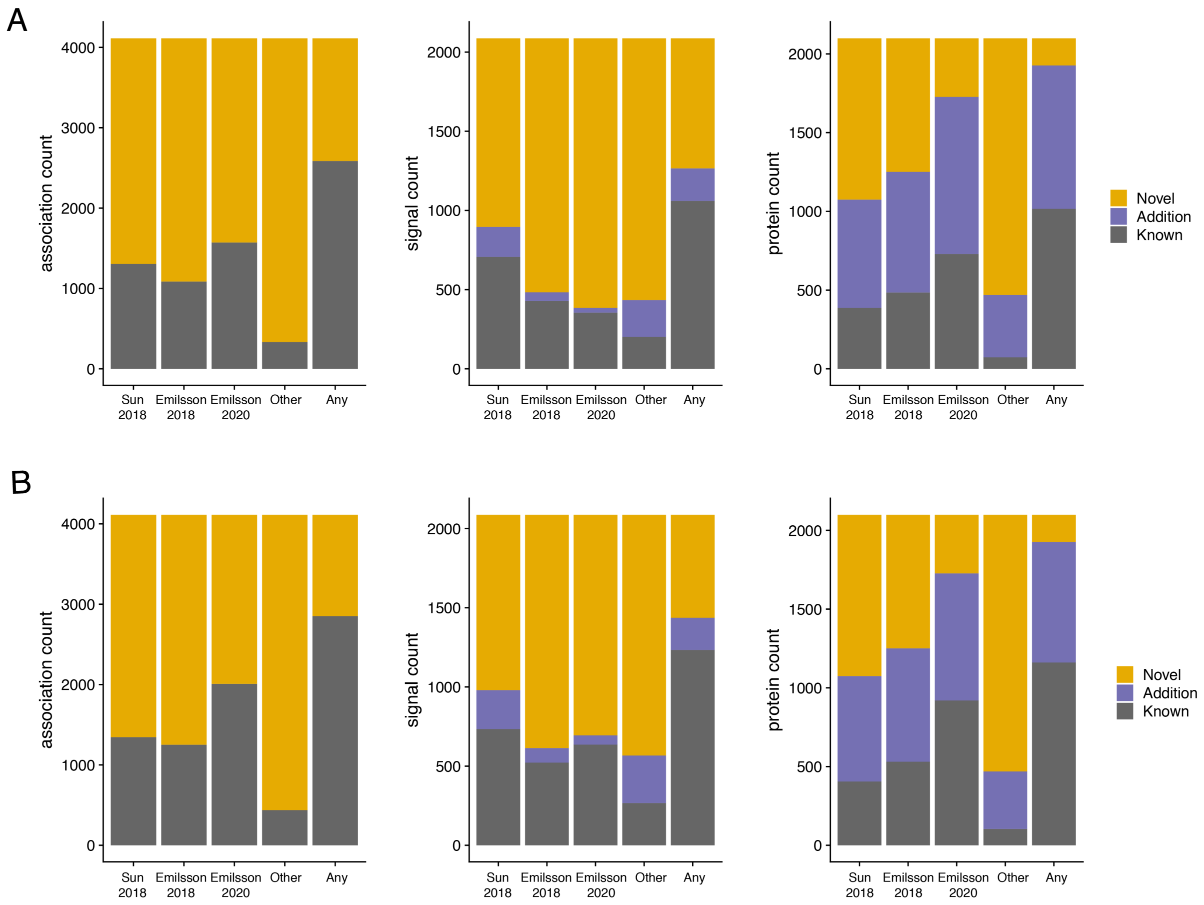

**Supplementary Figure 4** Comparison of independent study-wide significant pQTLs identified in the current study (n = 5,368) to those previously reported, using A) the primary LD threshold $r^2 > 0.9$ to define shared signals, and B) a more stringent LD threshold $r^2 > 0.5$ to define shared signals. The barplots show if a matching pQTL association (left panel) has been previously been reported, if a genetic signal (middle panel) has been reported to associate with the same (known) or another (addition) protein, and if the same (known) or another (addition) genetic signal has previously been reported for a given protein (right panel). See Supplementary Note Table 1 for exact numbers. An overview of the studies used for comparison is shown in Supplementary Table 5.

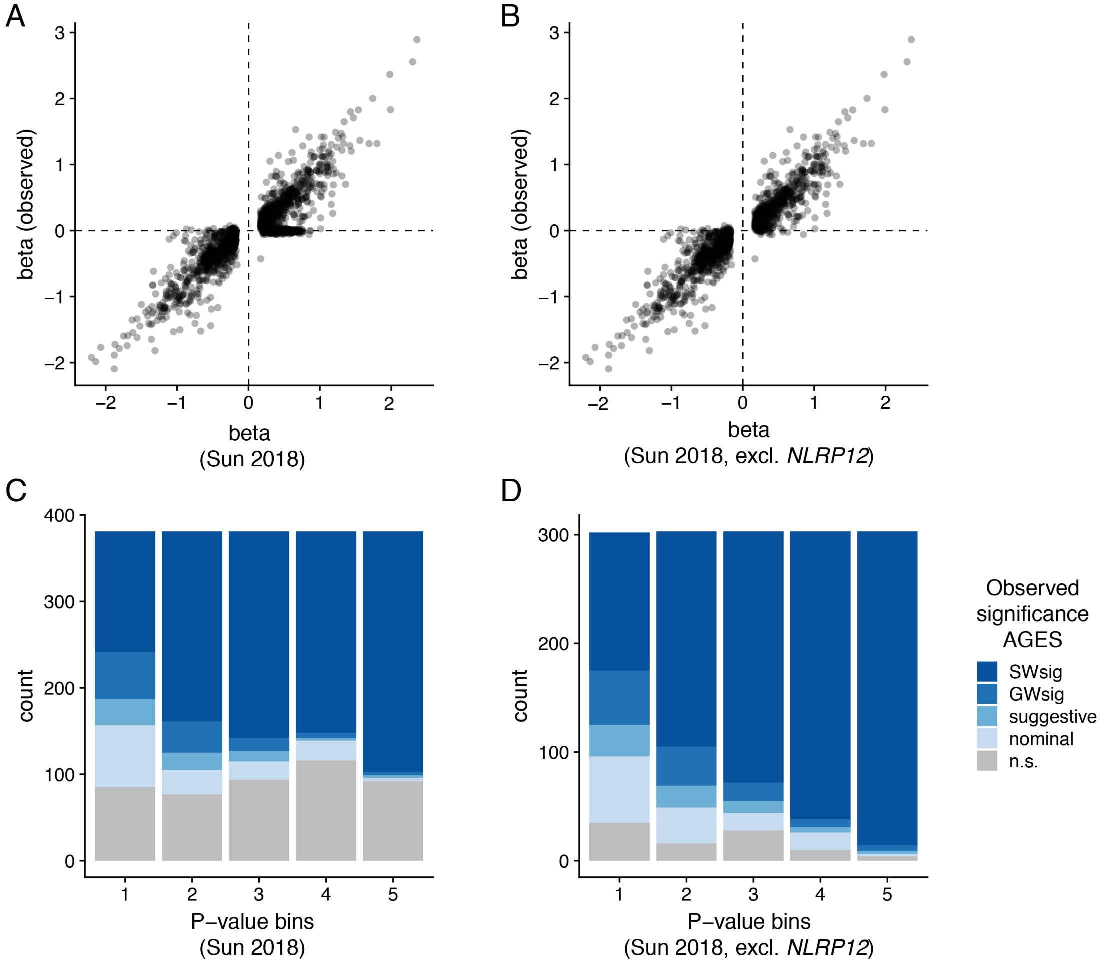

**Supplementary Figure 3** Replication of previously reported pQTLs from the INTERVAL study (Sun et al. 2018) in AGES. A-B) Comparison of reported pQTL beta values for (A) all independent study-wide significant pQTLs reported by Sun et al 2018 (B) all independent study-wide significant pQTLs reported by Sun et al 2018 but excluding the *NLRP12* locus, with matching pQTL beta values observed in the current study. C-D) Number of associations (pQTLs) colored by observed significance level for pQTLs in AGES and shown by significance bins based on reported pQTL P-values for (C) all independent study-wide significant pQTLs reported by Sun et al 2018 and (D) all independent study-wide significant pQTLs reported by Sun et al 2018 but excluding the *NLRP12* locus, where bin 5 has the lowest (most significant) reported P-values. SWsig: study-wide significant, $P<5\times10^{-8}/4782$, GWsig: genome-wide significant, $P<5\times10^{-8}$, suggestive: $P<1\times10^{-5}$, nominal: $P<0.05$, n.s.: not significant, $P>0.05$.

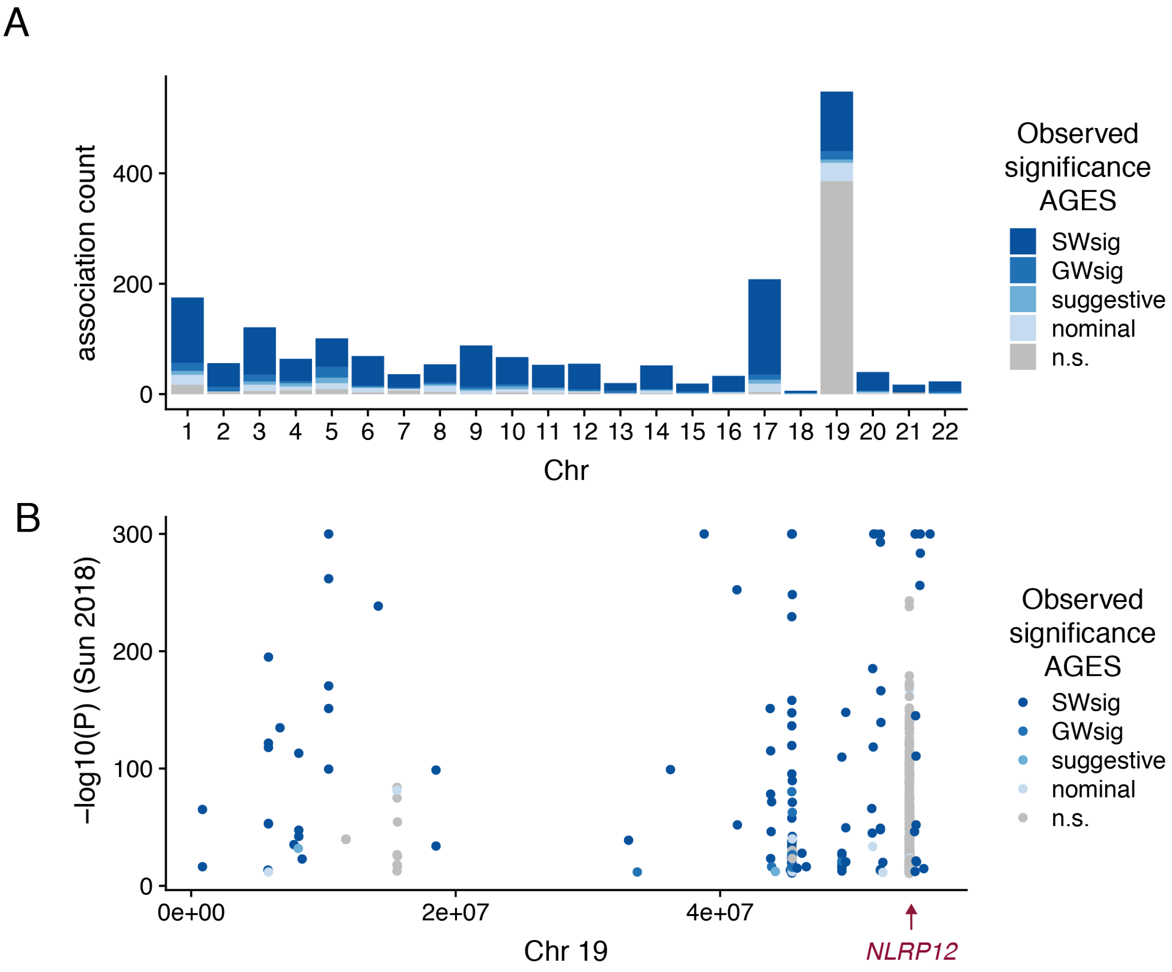

**Supplementary Figure 4** A) Number of reported independent study-wide significant associations (pQTLs) by Sun et al. 2018 from the INTERVAL study by chromosome, colored by observed significance level in AGES (n = 5,368). B) Independent study-wide significant pQTLs reported by Sun et al. 2018 on chromosome 19 colored by significance level in AGES (n = 5,368). The *NLRP12* locus is highlighted. SWsig: study-wide significant, $P<5\times10^{-8}/4782$, GWsig: genome-wide significant, $P<5\times10^{-8}$, suggestive: $P<1\times10^{-5}$, nominal: $P<0.05$, n.s.: not significant, $P>0.05$.

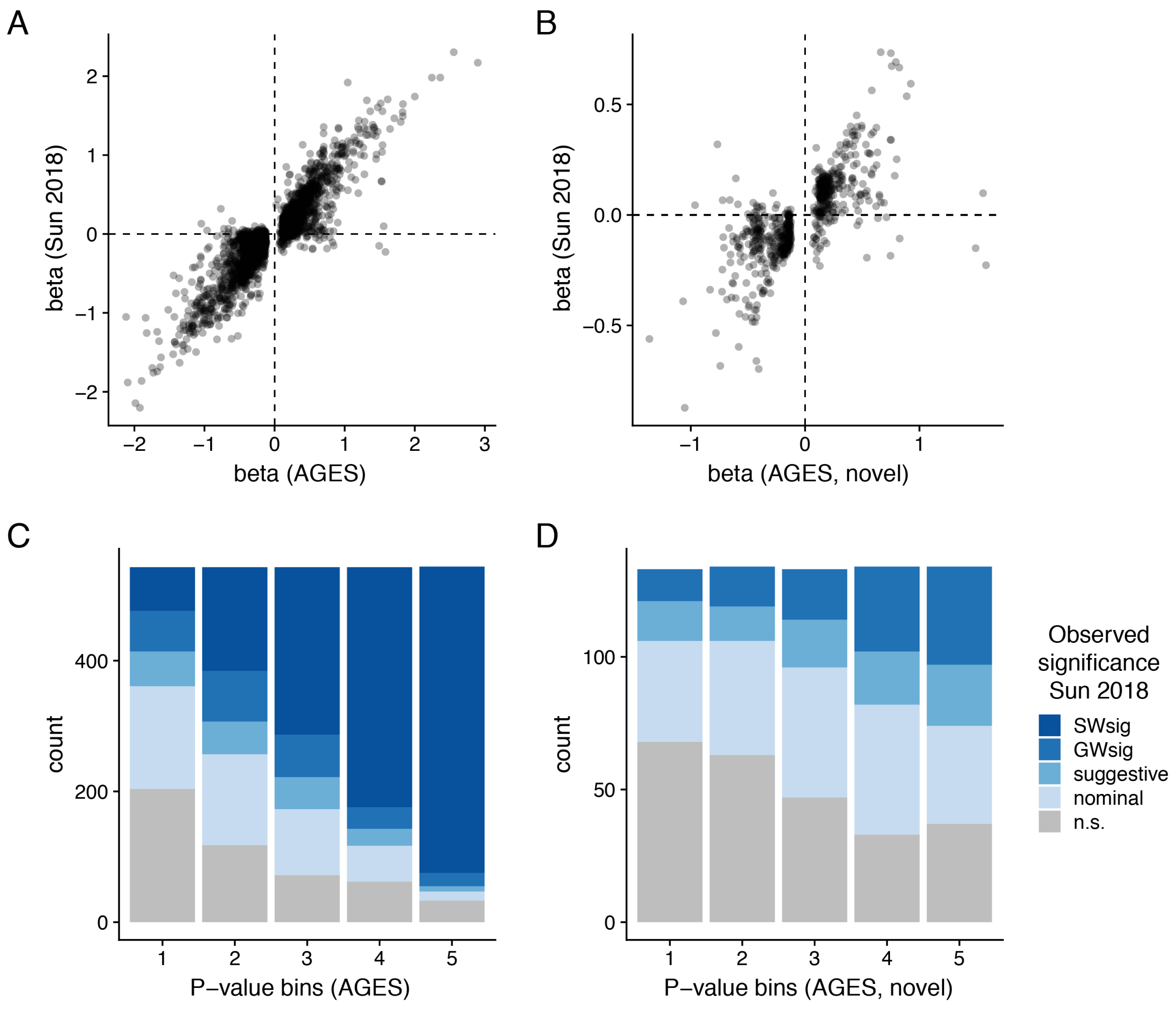

**Supplementary Figure 7** Replication of pQTLs identified in the current study using summary statistics from the INTERVAL study (Sun et al. 2018). A-B) Comparison of beta values for (A) all independent study-wide significant pQTLs in AGES and (B) independent study-wide significant pQTLs defined as novel in the current study, with matching pQTL beta values from Sun et al 2018. C-D) Number of associations (pQTLs) binned by P-values in the current study, shown for (C) all independent study-wide significant pQTLs and (D) independent study-wide significant pQTLs defined as novel in the current study, where bin 5 has the lowest (most significant) P-values, colored by significance level in the INTERVAL study (Sun et al. 2018). SWsig: study-wide significant, $P<5\times10^{-8}/3{,}283$ (number of SOMAmers in INTERVAL study), GWsig: genome-wide significant, $P<5\times10^{-8}$, suggestive: $P<1\times10^{-5}$, nominal: $P<0.05$, n.s.: not significant, $P>0.05$.

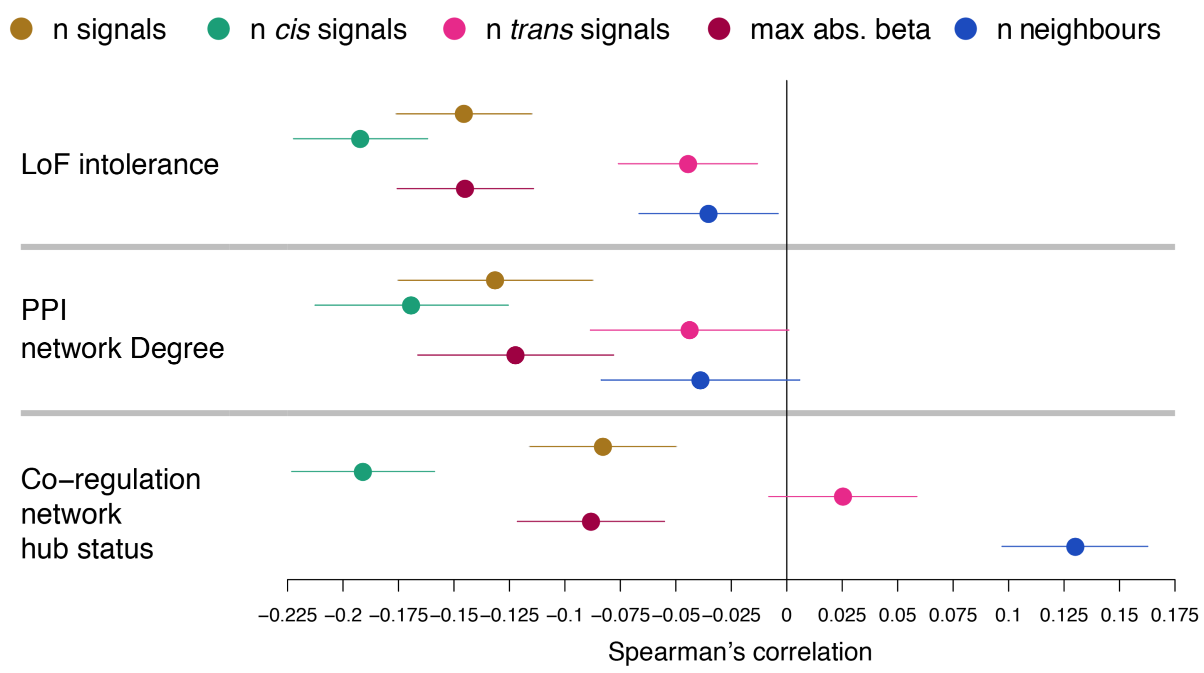

**Supplementary Figure 5** Spearman's correlation with 95% confidence intervals comparing genetic association profile characteristics (number of signals in total, cis or trans, largest absolute effect size among study-wide significant independent pQTLs, and number of neighbours defined as the number of proteins that share the same genetic signal) for proteins to metrics of loss-of-function (LoF) intolerance, protein-protein interaction (PPI) network degree and co-regulation network hub status.

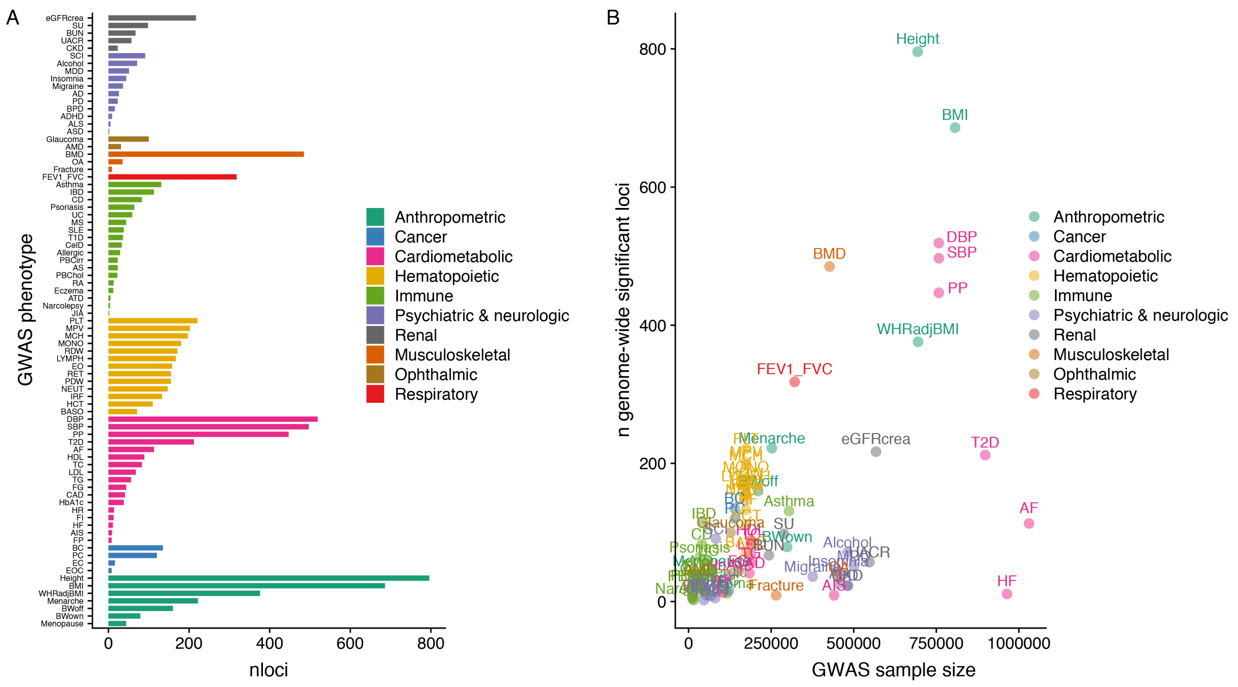

**Supplementary Figure 6** An overview of the GWAS sample sizes (x-axis) and the number of genome-wide significant loci (y-axis) for the 81 clinical traits and diseases for which GWAS summary statistics were included in the colocalization analysis with serum proteins. In particular, GWAS for immune traits tend to have smaller sample sizes, and subsequently fewer identified loci, than other trait categories.

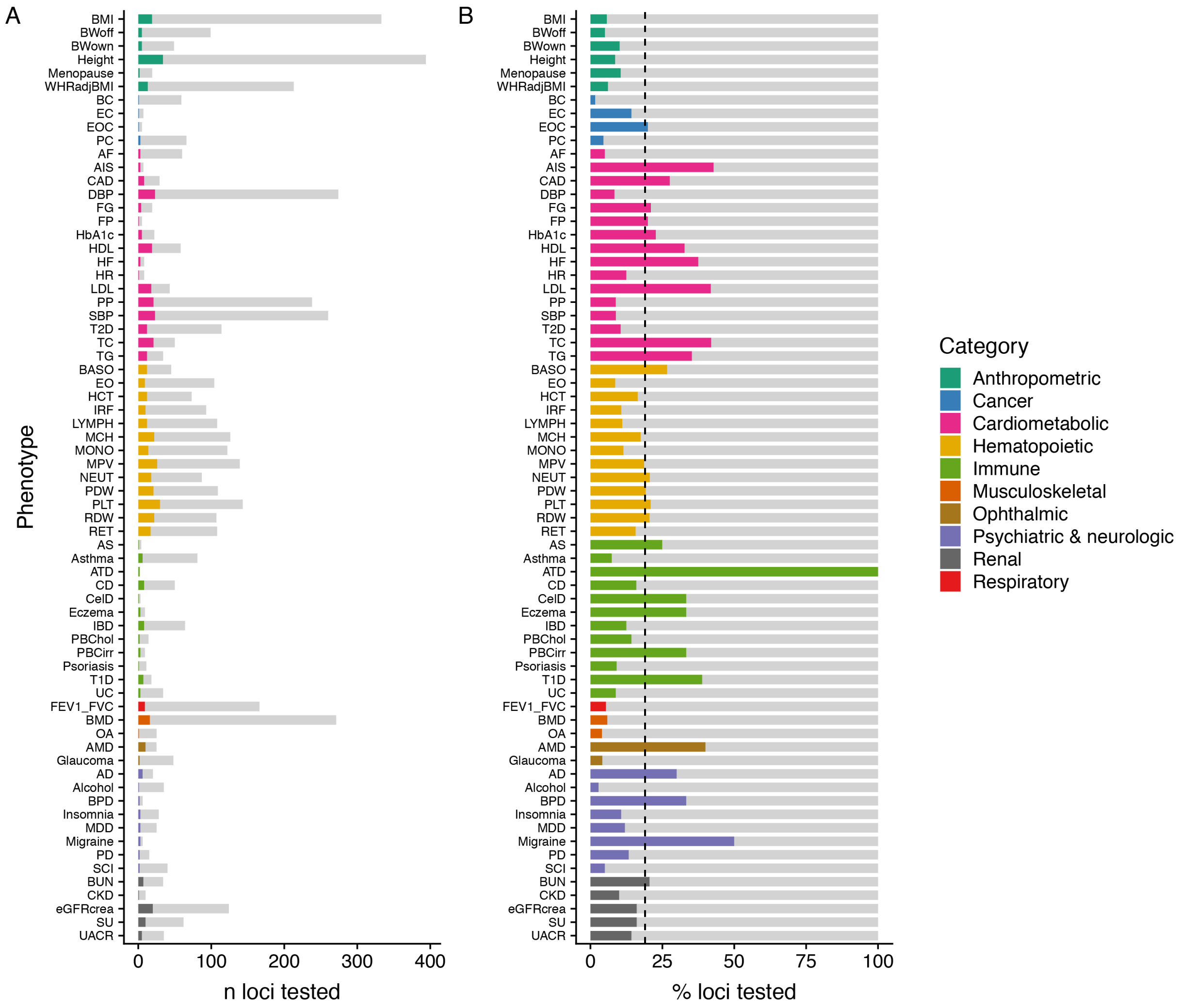

**Supplementary Figure 7** An overview of the A) number and B) proportion of GWAS loci that colocalize with any serum protein signal in the current study, restricted to the loci that were in the vicinity of a pQTL and thus tested. The dashed line indicates the average for the tested loci, or 19% that colocalize with at least one serum protein (whereas it is 11% when considering all GWAS loci).

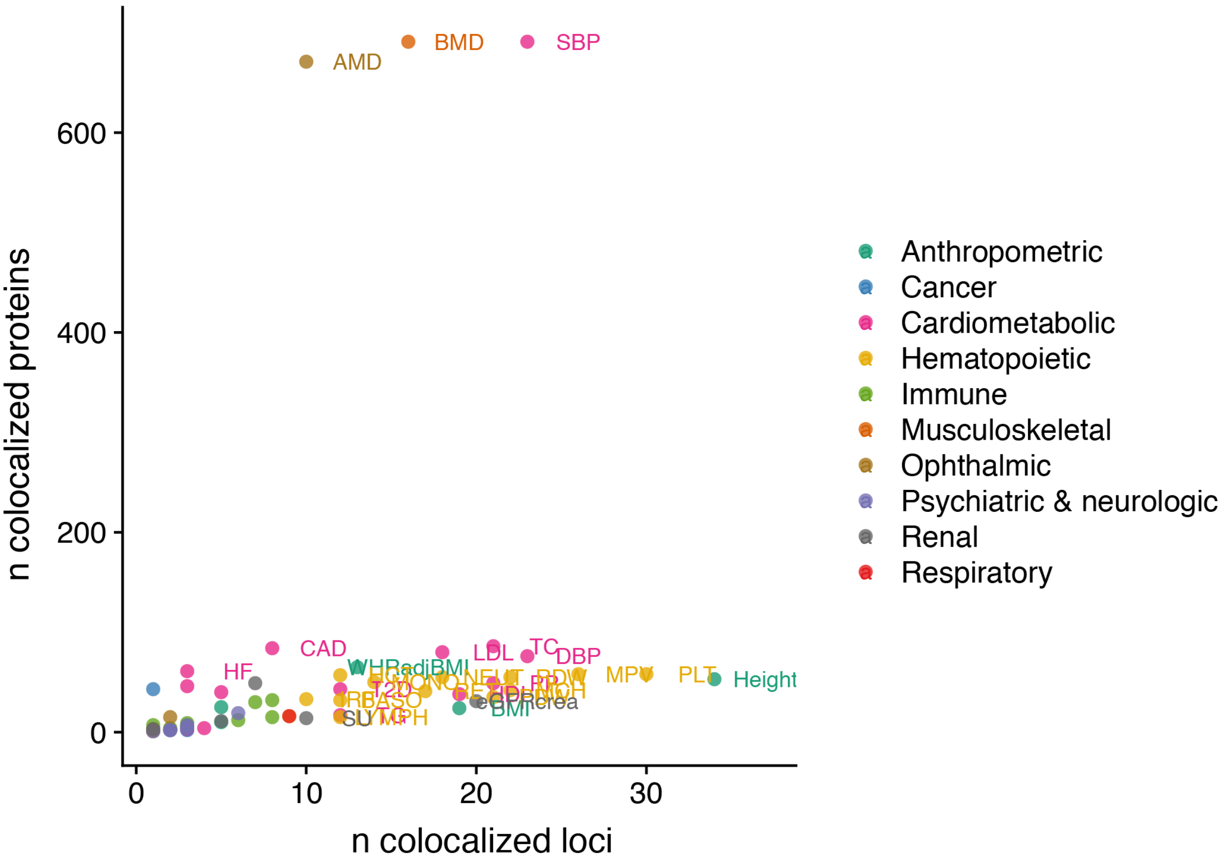

**Supplementary Figure 8** An overview of the number of colocalized proteins (y-axis) compared to the number of colocalized loci (x-axis) per trait. The three traits (AMD, BMD, SBP) at the top all colocalize with the *VTN* locus, which regulates 597 proteins.

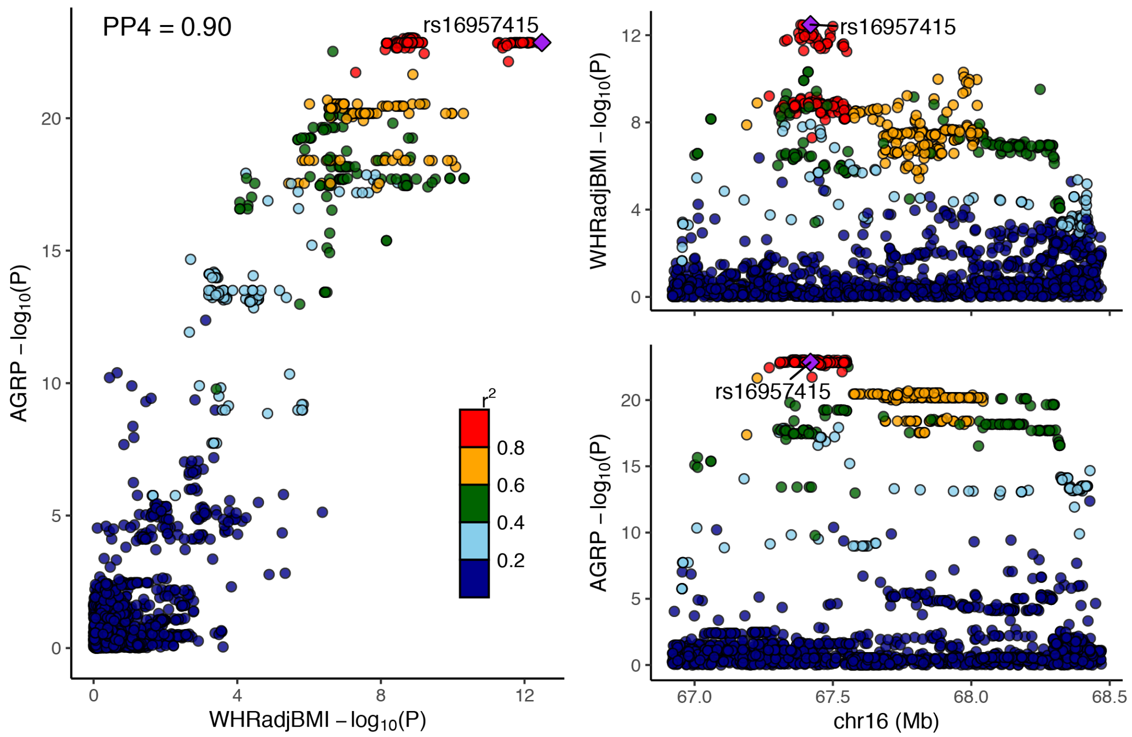

**Supplementary Figure 12** Colocalization of genetic associations for serum levels of the AGRP protein and waist-to-hip ratio adjusted for BMI (WHRadjBMI) on chromosome 16.

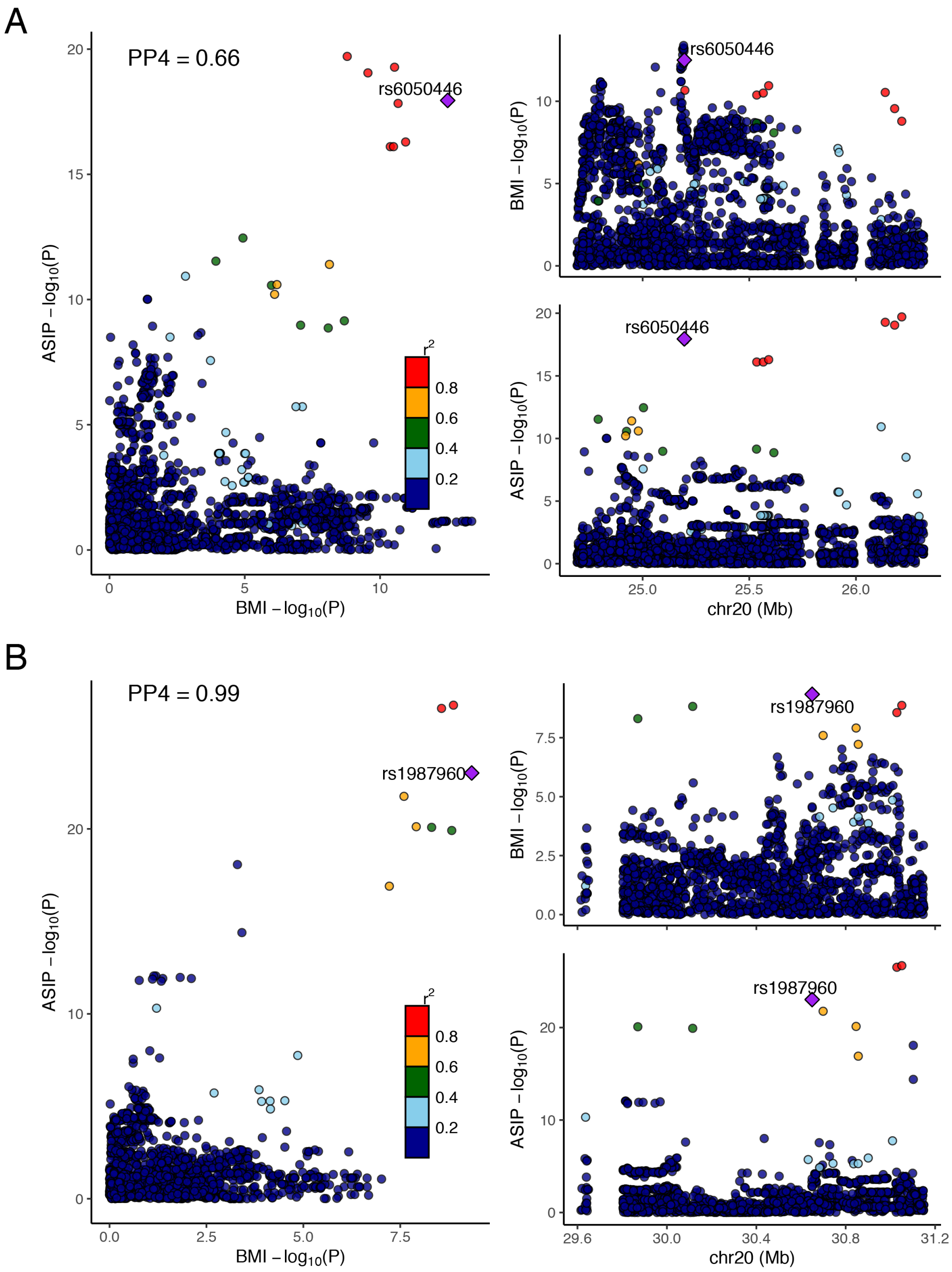

**Supplementary Figure 13** Colocalization of two genetic associations for serum levels of the ASIP protein and BMI on chromosome 20 at around A) 25.5Mb and B) 30.4Mb.

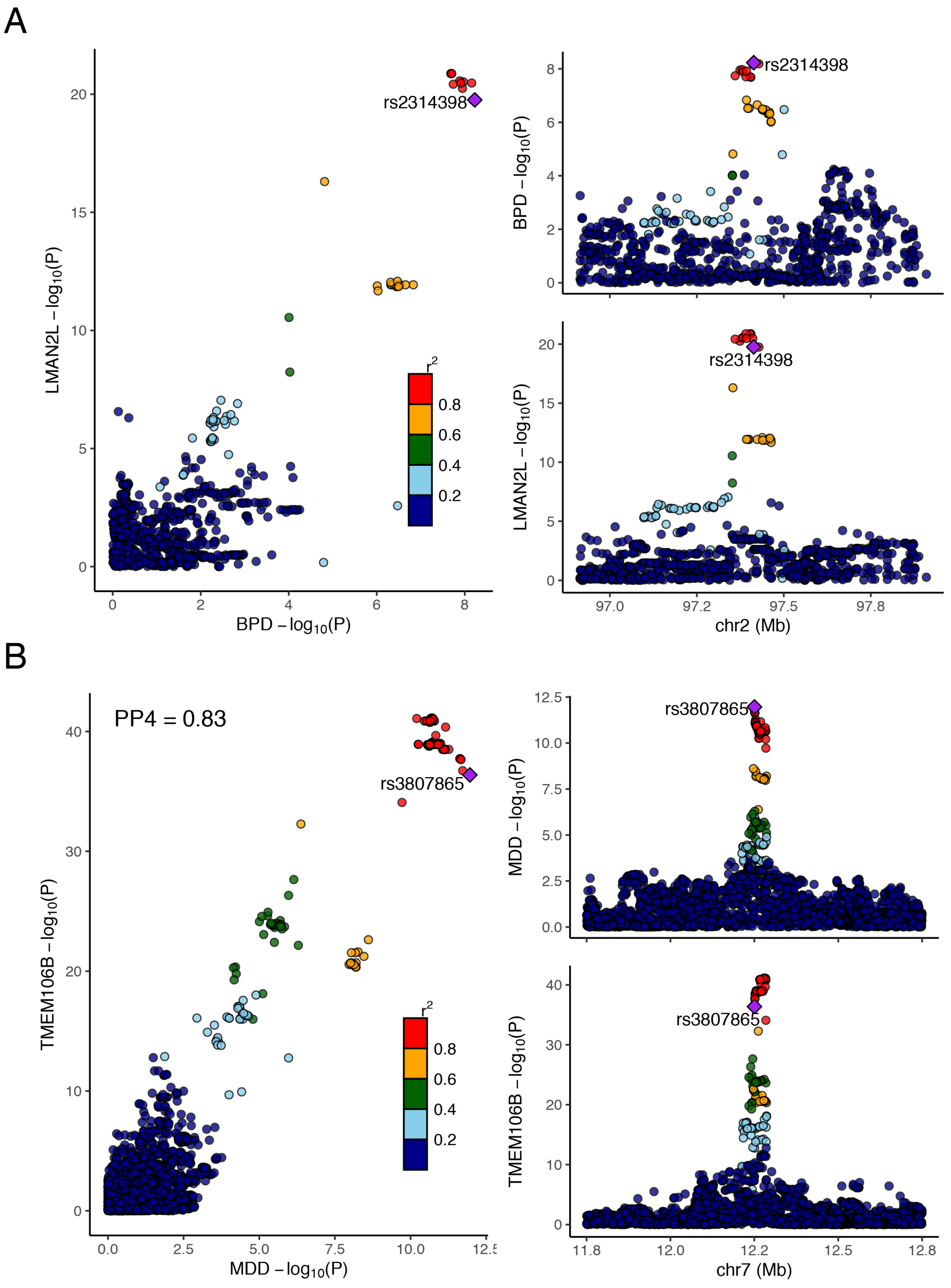

**Supplementary Figure 14** Colocalization between genetic associations for A) serum levels of the LMAN2L protein and bipolar disorder (BPD) on chromosome 2 and B) serum levels of the TMEM106B protein and major depressive disorder (MDD) on chromosome 7.

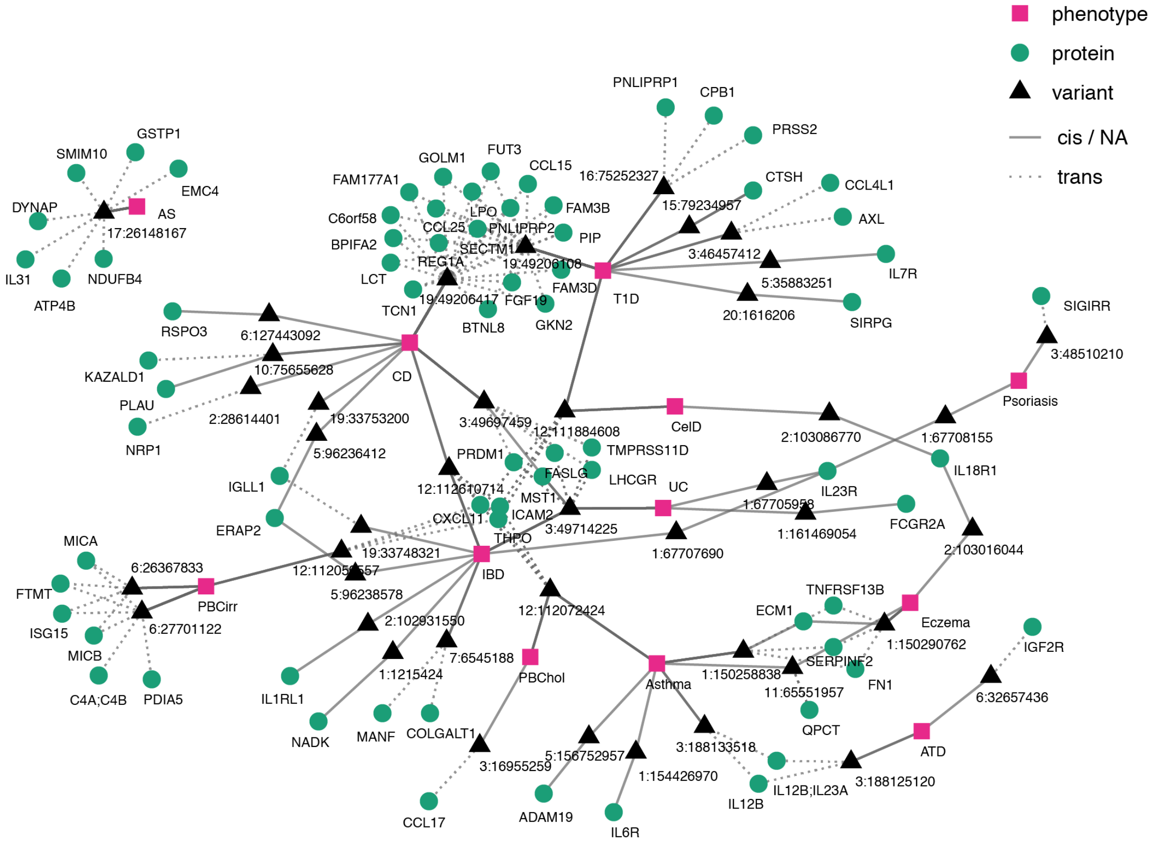

**Supplementary Figure 15** Immune disease colocalization network, illustrating how diseases (pink squares) are linked through the overlap of colocalized proteins (green circles). Diseases and proteins are linked through lead variants (black triangles) in the loci where genetic signals colocalize, where the pQTL may be in *cis* (solid line) or *trans* (dotted line).

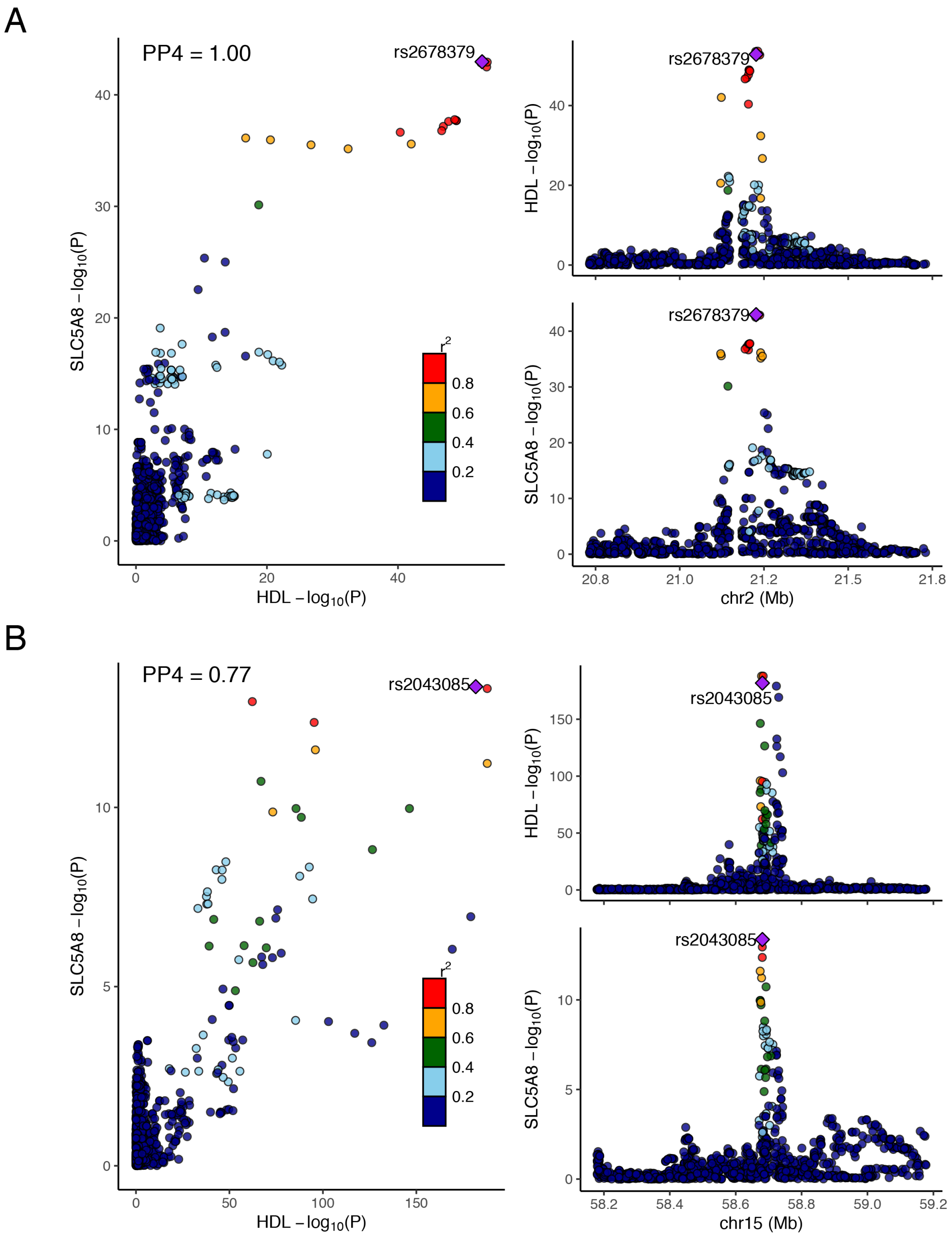

**Supplementary Figure 16** Colocalization between genetic associations for serum levels of the SLC58A protein and HDL cholesterol on A) chromosome 2 and B) chromosome 15.

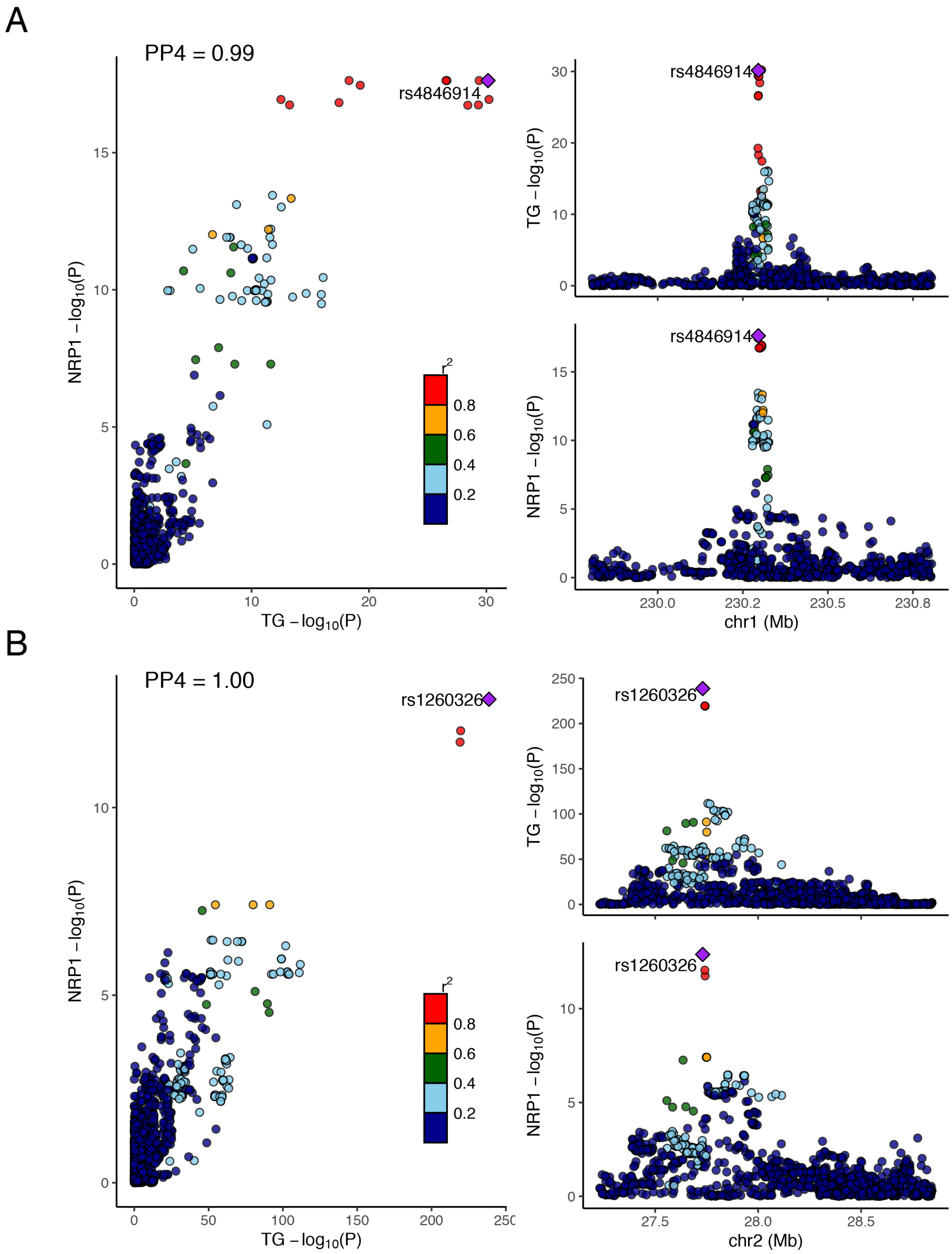

**Supplementary Figure 17** Colocalization between genetic associations for serum levels of the NRP1 protein and triglycerides (TG) on A) chromosome 1 and B) chromosome 2.

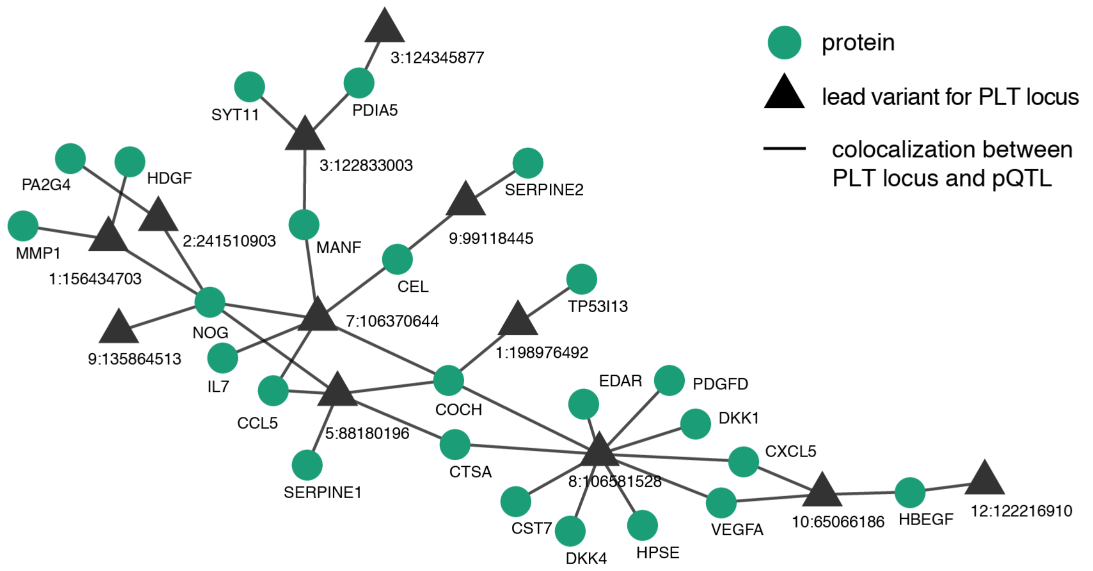

**Supplementary Figure 18** Platelet colocalization network, illustrating how many of the same proteins (green circles) colocalize with GWAS signals for platelet count (PLT) in multiple loci, represented by lead variants (black triangles). For example, pQTLs for NOG and COCH colocalize with GWAS signals for platelet counts in five and four loci, respectively.

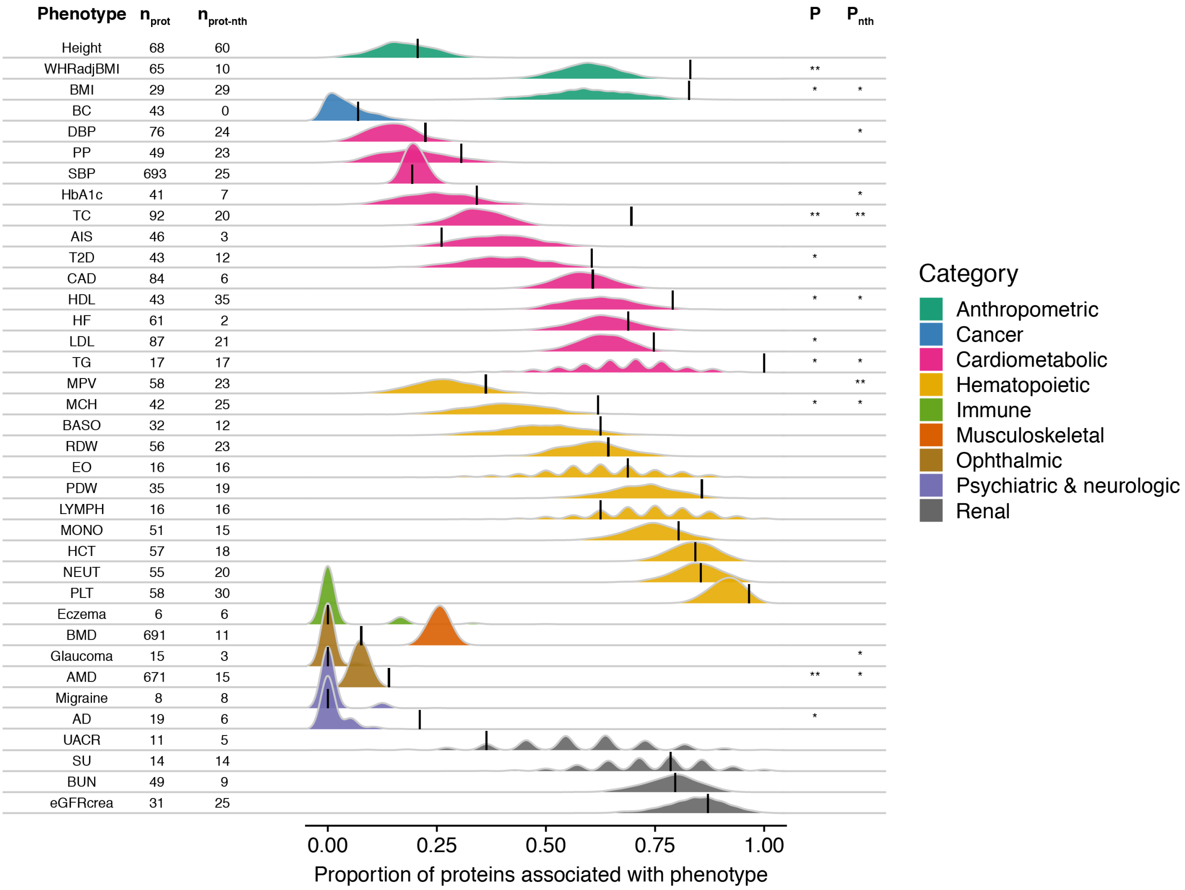

**Supplementary Figure 19** Ridgeline plot illustrating for each GWAS phenotype the proportion of colocalized proteins (including only study-wide significant pQTLs) that were significantly (FDR<0.05) associated with the same trait in AGES (n = 5,457) (black lines) compared to 1000 randomly sampled sets of proteins of the same size (density curves). The number of colocalized proteins for each trait are provided on the left-hand side, along with the number of proteins remaining after the removal of proteins originating from loci with 5 or more colocalized proteins from the analysis, annotated as "no trans-hotspots" (nth). Empirical p-values for significant enrichment of protein-phenotype associations are denoted as such: *P < 0.05, **P < 0.001.

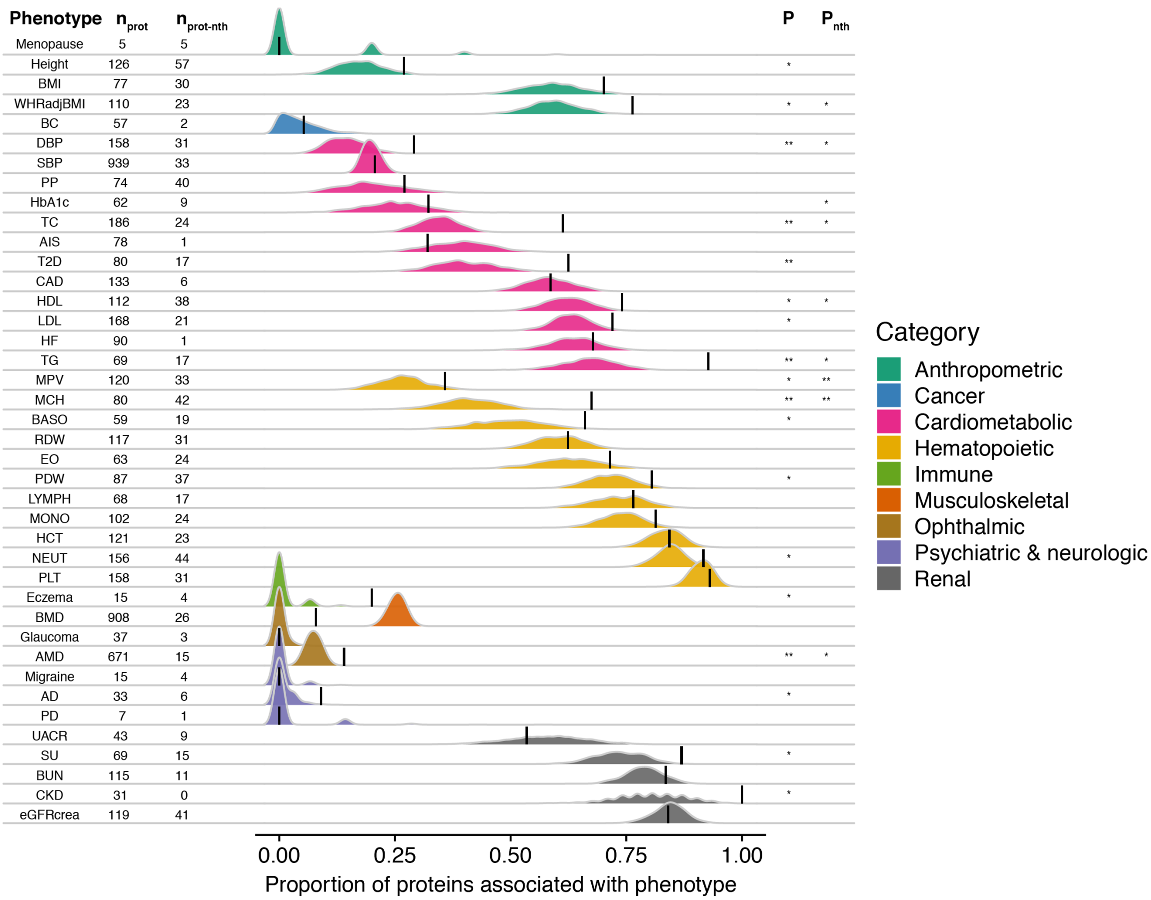

**Supplementary Figure 20** Ridgeline plot illustrating for each GWAS phenotype the proportion of colocalized proteins (including all genome-wide significant ($P<5\times10^{-8}$) pQTLs) that that were significantly (FDR<0.05) associated with the same trait in AGES (n = 5,457) (black lines) compared to 1000 randomly sampled sets of proteins of the same size (density curves). The number of colocalized proteins for each trait are provided on the left-hand side, along with the number of proteins remaining after the removal of proteins originating from loci with 5 or more colocalized proteins from the analysis, annotated as “no trans-hotspots” (nth). Empirical p-values for significant enrichment of protein-phenotype associations are denoted as such: *P < 0.05, **P < 0.001.
